# *In vivo* CRISPR screens identify GRA12 as a transcendent secreted virulence factor across *Toxoplasma gondii* strains and mouse subspecies

**DOI:** 10.1101/2024.09.10.611481

**Authors:** Francesca Torelli, Simon Butterworth, Eloise Lockyer, Ok-Ryul Song, Jennifer Pearson-Farr, Moritz Treeck

## Abstract

*Toxoplasma gondii* parasites exhibit extraordinary host promiscuity owing to over 250 putative secreted proteins that disrupt host cell functions, enabling parasite persistence. However, most of the known effector proteins are specific to *Toxoplasma* genotypes or hosts. To identify virulence factors that function across different parasite isolates and mouse strains that differ in susceptibility to infection, we performed systematic pooled *in vivo* CRISPR-Cas9 screens targeting the *Toxoplasma* secretome. We identified several proteins required for infection across parasite strains and mouse species, of which the dense granule protein 12 (GRA12) emerged as the most important effector protein during acute infection. GRA12 deletion in IFNγ-activated macrophages results in collapsed parasitophorous vacuoles and increased host cell necrosis, which is partially rescued by inhibiting early parasite egress. GRA12 orthologues from related coccidian parasites, including *Neospora caninum* and *Hammondia hammondi,* complement TgΔGRA12 *in vitro*, suggesting a common mechanism of protection from immune clearance by their hosts.

## Introduction

*Toxoplasma gondii* is one of the most prevalent parasites worldwide and infects any nucleated cell of warm-blooded animals, including humans ^1^. Most isolates can be grouped into three lineages – type I, II and III – with progressively decreasing virulence in mice ^2^. Less virulent type II and III strains dominate in Europe and North America ^3^, and infection is generally asymptomatic in humans. Less common but more virulent type I strains have been associated with clinical complications in humans, such as visual impairment and foetal malformations ^4,5^. In South America, particularly in Brazil, a multitude of genetically diverse isolates have been identified over the last decades^6^. These so-called “atypical” isolates ^7^ caused several lethal outbreaks affecting immunocompetent individuals, high miscarriage rates and lack of cross-protective immunity ^8,9^. While work of the last decades largely focussed on identifying virulence factors that make one strain more virulent than another, the factors conserved across strains and conferring *Toxoplasma* success in a wide host range remain unknown to date.

*Toxoplasma*’s capacity to escape immune clearance is largely conferred by a pool of proteins secreted from two secretory organelles during and after invasion: rhoptries (ROPs) and dense granules (GRAs). Secreted proteins generate a supporting environment for *Toxoplasma* growth by rewiring host cell transcription and interfering with the host cell immune machinery ^10^. For example, the dense granule inhibitor of STAT1 transcriptional activity (IST) blocks the upregulation of interferon-stimulated genes in naïve cells and subsequent parasite clearance ^11–13^, while the rhoptry kinase ROP16 drives macrophage polarisation to support parasite growth ^14,15^. Secreted proteins either reside within the non-fusogenic parasitophorous vacuole (PV) created by the actively growing intracellular tachyzoites, span the PV membrane (PVM) or cross the PVM into the host cytoplasm. Recently, the number of secreted proteins has been estimated to total around 250, the vast majority without functional annotations ^16^.

To date, the best characterised virulence factor is probably the serine-threonine kinase ROP18, which is directly associated with parasite virulence ^17,18^. ROP18, together with the pseudokinase ROP5, inactivates the main murine defence mechanism against intracellular pathogens: the Immunity-Related GTPases (IRGs) ^17,19^. The IRG family has specifically expanded in the murine genome ^20^, where it includes 23 members, and has been associated with host resistance to intracellular pathogens of parasitic, bacterial and fungal origin ^21–23^. Specifically, IRGa6, IRGb6, IRGd and IRGb10, are pivotal for host survival to *Toxoplasma* infection via coordinated loading on the PVM ^24^, which depends on the regulatory IRGs, namely IRGm1 and IRGm3 ^25,26^. Irgm1/m3 are also paramount for the Ubiquitin-mediated degradation of the *Toxoplasma* vacuole and for general cellular homeostasis ^27,28^. In human cells, IRGs are almost entirely missing. Another family of GTPases, the guanylate binding proteins (GBPs), is involved in *Toxoplasma* restriction in both human and murine hosts. mGBP1 and mGBP2 control *Toxoplasma* infection in murine cells ^29,30^, and hGBP1, hGBP2 and hGBP5 in human macrophages ^31^. Recent work showed that murine resistance to *Toxoplasma* relies on the PV nitrosylation and collapse of GBP2-recruited vacuoles^32^. IRGs and GBPs loading on the PVM results in membrane ruffling, vacuolar rounding and breakage eventually resulting in parasite elimination via an unknown mechanism ^24,29,30^. Parasite clearance leads to host cell death, which is considered a hallmark of host resistance to infection ^33,34^. The activation of specific programmed host cell death pathways, like apoptosis and pyroptosis ^35,36^, were observed following loading of IRGs and GBPs.

All main mouse subspecies – i.e. *M. m. domesticus*, *M. m. musculus* and *M. m. castaneus* – can control infection with the less virulent type II and III Toxoplasma strains, however the outcomes after infection with the highly virulent type I parasites are different. For example, infection with type I strains is lethal in the most commonly used *M. m. domesticus* mouse strains, e.g. C57BL/6J, where ROP18 inhibits the vacuolar accumulation of host IRGa6 ^19^. Type II and III parasites express less virulent isoforms or undetectable levels of ROP18 respectively ^37^. Due to polymorphisms in the IRG loci, *M. m. castaneus* and *M. m. musculus* strains are resistant to ROP18 function and survive infection ^33,38,39^. Therefore, ROP18 is a virulence factor, but only in some murine subspecies and parasite strains. Most South American *Toxoplasma* strains are lethal in all three subspecies ^38,39^, suggesting the presence of unknown parasite virulence factors in this *Toxoplasma*-host combination.

Most virulence factors characterised to date are specific to certain clonal strains, host species or cell types, or a combination of them ^40^. This is likely biased by the initial studies to identify strain-specific virulence factors based on *Toxoplasma* genetic crosses, and by the predominant use of *M. m. domesticus* laboratory mouse strains as model organisms ^41^. This is of particular importance when considering the key role of rodents in *Toxoplasma* transmission, as they are major prey of felines, the definitive host of the parasite. Thus, whilst strain- and species-specific virulence factors have been identified, we do not know which virulence factors are important in all parasite strains to colonise different hosts. If they exist, these conserved effectors significantly contribute to *Toxoplasma* promiscuity and abundance in nature.

To identify common virulence factors acting across mouse strains of different genetic backgrounds, we performed targeted CRISPR-Cas9 screens in three *Toxoplasma* strains belonging to the common lineages and the atypical VAND strain, in different murine subspecies. The dense granule protein 12 (GRA12) emerged as the most important secreted protein for survival in the mouse peritoneum during acute infection and in IFNγ-activated macrophages, irrespective of parasite or mouse genetic backgrounds. Deletion of GRA12 led to increased regulated necrotic host cell death, partially caused by parasite early egress. A collapsed vacuolar space indicated a function of GRA12 in maintaining this replicative niche during the IFNγ-mediated host cell defence. Complementation of GRA12 deletion with GRA12 orthologues from two closely related parasite species suggested that the protective mechanism is conserved and important beyond *Toxoplasma* infections.

## Results

### Targeted *in vivo* CRISPR screens identify GRA12 as a *Toxoplasma* strain-transcendent secreted virulence factor in the mouse peritoneum

To identify virulence factors that function across parasite strains and mouse species, we generated pooled CRISPR mutant libraries as previously described ^42–44^ in RH ΔHXGPRT and PRU ΔHXGPRT (hereafter named RH and PRU), VEG and VAND parasites to encompass type I, type II, type III and atypical strains, respectively. Parasite knock-out (KO) pools for 253 predicted rhoptry and dense granule proteins were injected intraperitoneally (i.p.) into five mice per parasite strain (Figure 1A). To avoid skewing the results towards the known strain-specific virulence factor ROP18 in susceptible *M. m. domesticus* mice, we used selected mouse subspecies for the CRISPR screens: for the more virulent VAND and RH strains we used the *M. m. castaneus* CAST/Eij and *M. m. musculus* PWD/Phj, respectively (called “IRG resistant mice”). The different subspecies were used in these two screens due to loss of mouse colonies during the COVID-19 pandemic. The PRU and VEG screens were performed in *M. m. domesticus* C57BL/6J (called “IRG susceptible mice”). Five days post infection, parasites were recovered from the peritoneal exudates, their sgRNAs amplified by PCR and sequenced to determine their relative abundance before and after *in vivo* selection (Figure 1A). Genes that contribute to survival *in vivo* will display a negative log_2_ fold change (L2FC), because parasites lacking those genes will be cleared during the infection.

**Figure 1.**
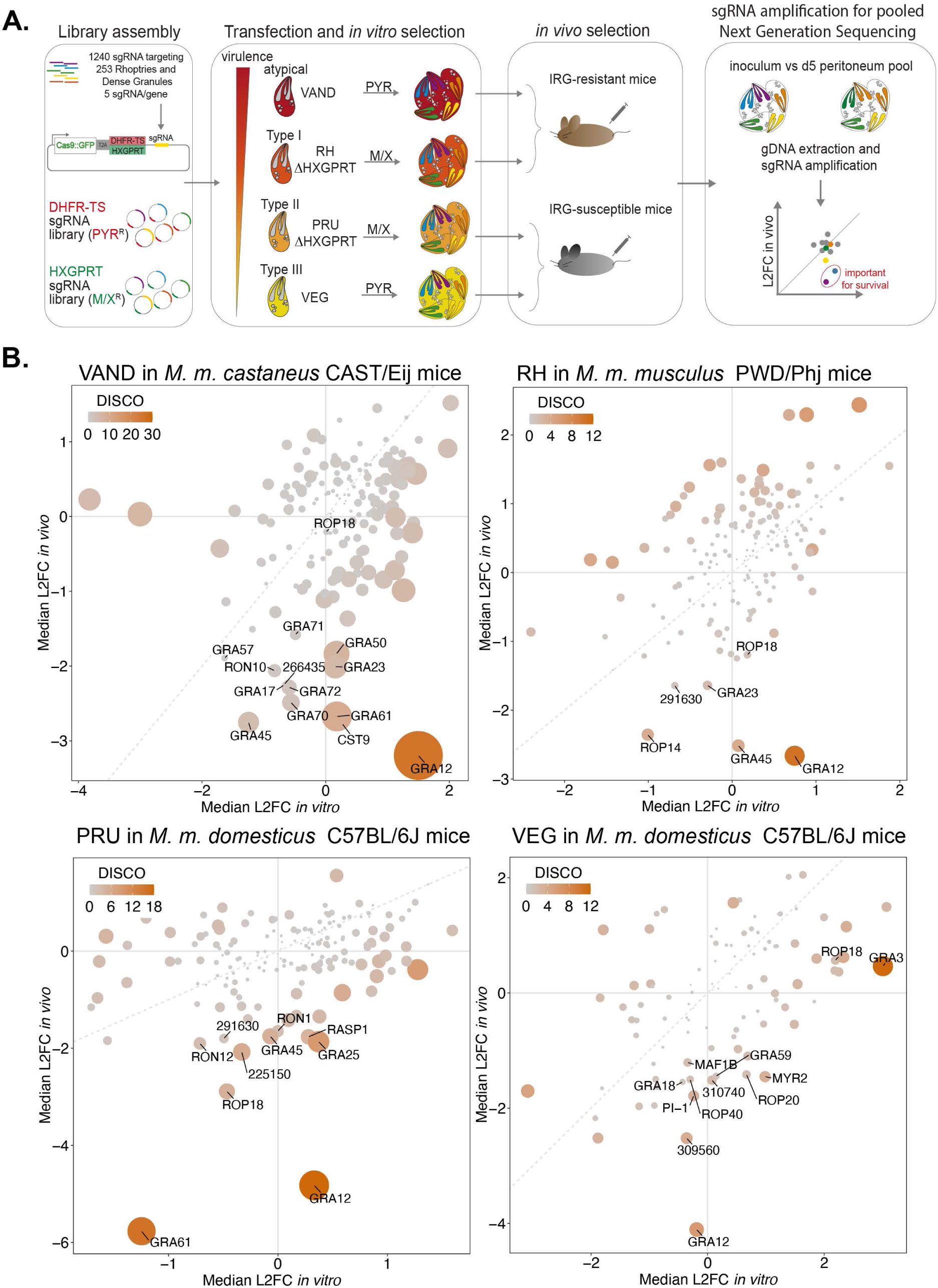
CRISPR screens *in vivo* identify GRA12 as the most important common secreted virulence factor for survival across clonal and atypical *Toxoplasma* strains. (A) Scheme of the CRISPR screen pipeline. Two protospacers libraries targeting the putative *Toxoplasma* secretome were assembled and included either the DHFR-TS or the HXGPRT selection cassette to create pyrimethamine-resistant (PYR^R^) or mycophenolic acid/xanthine-resistant (M/X^R^) parasite KO pools. The wild-type strains VAND and VEG were transfected with the DHFR-TS library, while the ΔHXGPRT strains RH and PRU were transfected with the HXGPRT library. The resulting KO parasite pools were injected in the peritoneum of 5 mice/parasite strain, retrieved after 5 days and expanded for one lytic cycle *in vitro* before gDNA extraction and sgRNA amplification for sequencing. The relative abundance of each guide (L2FC) at day 5 versus inoculum indicates the importance for *in vivo* survival of the relative *Toxoplasma* gene. (B) Scatter plots of the median L2FC for each gene *in vitro* and *in vivo* in CRISPR screens of, in order from the top left: VAND in CAST/EiJ mice, RH ΔHXGPRT in PWD/Phj mice, PRU ΔHXGPRT and VEG in C57BL/6J mice. The colour and size of each point reflects the Discordance/Concordance (DISCO) score, and the dashed grey line indicates equal L2FC. Displayed data are thresholded for L2FC *in vitro* >-1 according to published *in vitro* whole genome screens ^46^. Data points with a L2FC *in vivo* <-1.5 are labelled in addition to ROP18 as control.

We observed comparable variability, in terms of median absolute deviation (MAD) of the L2FC and sgRNA loss, *in vitro* and *in vivo* in three of the four screens. The VEG screen displayed a higher variability (Figure S1A-D), which is probably promoted by the strong tendency of VEG to encyst ^45^. Encystation leads to growth arrest, which likely increases the dropout rate of mutants from the pool and reduction of valid data points. Regardless, *in vitro* and *in vivo* L2FC scores correlate well with previous screens performed in our laboratory in the PRU strain (Figure S1E-F). Data points with a more pronounced difference between the median L2FC *in vivo* and *in vitro* display a higher discordance/concordance (DISCO) score which is adjusted for the p-value of each parameter (Figure 1B). All raw and normalised read counts, L2FC and DISCO scores for each screen are reported in Table S2-5. To identify virulence factors common between strains, we ranked the candidates based on the difference between L2FC *in vivo* and *in vitro* (L2FC DIFF, Table S6) using the cumulative score for each candidate across all screens (Table 1). The relatively high loss of mutants in the VEG screen led to a loss of data points for most genes identified as important in the other *Toxoplasma* strains. However, relaxing the inclusion criteria from three guides per gene to one guide per gene for the VEG screen resulted in data points for all ranked virulence factors (Figure S1G and Table 1). While the values from the VEG screen should be taken with some caution, they indicate that several proteins play important roles across strains.

**Table 1.** The five top common virulence factors across clonal and atypical *Toxoplasma* strains. Genes were ranked based on the difference between the median L2FC *in vivo* and *in vitro* across all strains. The median L2FC *in vivo* and the DISCO score are reported for the five top hits. For the VEG screen numbers are reported also from the quantification with relaxed selection (>=1 guide/ gene). Values obtained only with relaxed selection are indicated with an asterisk, with the number of guides used for quantification in brackets.

| Rank | Gene | Code | VAND |  | RH |  | PRU |  | VEG |  |
| --- | --- | --- | --- | --- | --- | --- | --- | --- | --- | --- |
|  |  |  | L2FC *in vivo* | DISCO | L2FC *in vivo* | DISCO | L2FC *in vivo* | DISCO | L2FC *in vivo* | DISCO |
| 1 | GRA12 | _288650 | -3.20 | 26.04 | -2.66 | 10.73 | -4.82 | 17.71 | -4.11 | 5.87 |
| 2 | GRA45 | _316250 | -2.75 | 4.55 | -2.52 | 4.01 | -1.75 | 4.89 | -5.25 (1)* | 5.25* |
| 3 | GRA61 | _269950 | -2.67 | 9.52 | -0.41 | 0.36 | -5.77 | 15.77 | -3.70 (2)* | 4.11* |
| 4 | GRA50 | _203600 | -1.84 | 7.08 | 0.32 | 1.72 | -0.21 | 3.53 | -2.47 (2)* | 4.31* |
| 5 | GRA23 | _297880 | -2.01 | 5.66 | -1.64 | 2.10 | -0.16 | 0.42 | -3.93 (1)* | 8.50* |

The dense granule protein GRA12 is most strongly correlated with reduced fitness upon deletion across all screens, regardless of the mouse or parasite backgrounds (Figure 1B and Table 1). In contrast, and as expected, correlation between ROP18 deletion and parasite fitness depends on parasite and mouse genotypes. Another important protein identified across the different screens is GRA45, which is involved in the correct localisation of proteins in the PVM ^47^, one of which, GRA23, was also identified in our screens ^48^. Furthermore, GRA61 and GRA50 have been identified to be fitness conferring *in vivo* in all performed CRISPR screens to date ^42,49–51^, but their functions remain unknown.

The screens also suggest that novel strain-specific virulence factors exist, such as the predicted dense granule protein TGGT1_291630 orthologs in PRU and RH, or orthologs of the rhoptry proteins TGGT1_309560 and TGGT1_266435 in VEG and VAND respectively. Interestingly, three proteins (GRA57, GRA70 and GRA71) previously identified as important factors for *Toxoplasma* survival in human cells ^44,52^, are uniquely identified in the VAND screen, suggesting a strain-specific role of this complex in this parasite and mouse combination that deserves further investigation.

While we cannot exclude that we missed some virulence factors because of the variability of the VEG screen, GRA12 emerges as the most critical virulence factor between all tested parasite strains in all mouse genetic backgrounds. GRA12 has been previously identified in *in vivo* CRISPR screens performed in type I and II strains by our research group and others ^42,49–51^ and is important for the survival of IFNγ-mediated restriction *in vitro* ^53^ and *in vivo* ^50^. However, the importance of GRA12 in other strains, such as type III and atypical, and how it mediates *Toxoplasma* survival had not been previously characterised.

### GRA12 is a virulence factor in the South American VAND strain *in vivo*

To validate GRA12 as a virulence factor in IRG resistant mice, we generated a ΔGRA12 and a GRA12-complemented strain in the atypical South American VAND parasite strain. The endogenous *Gra12* locus was replaced with a mCherry expression cassette via CRISPR-Cas9 and loss of the GRA12 expression was validated by immunofluorescence detection with an anti-GRA12 antibody (Figure S2A-B). An HA-tagged complemented strain (GRA12::HA) was established via homologous recombination in the *Uprt* locus (Figure S2C). Western blot analysis confirmed a GRA12::HA band at the expected size of 48 kDa, and GRA12::HA was localised to the intravacuolar space by immunofluorescence (Figure 2A-B). In line with an *in vitro* fitness score of +1.75 ^46^, lack of GRA12 confers a modest growth advantage to *Toxoplasma* in human fibroblasts in plaque assays, which is lost in the complemented strain (Figure S2D). Intraperitoneal infection of IRG resistant PWD/Phj mice with a dose of 50 to 500 parental and GRA12-complemented parasites was lethal, while mice infected with up to hundred times higher doses of ΔGRA12 parasites survived (Figure 2C). The surviving mice presented with VAND ΔGRA12 cysts in the brain, showing that GRA12 is not essential for latent stage formation *in vivo* (Figure 2D). However, as wildtype (WT)-infected mice succumb to infection before cyst formation, we cannot draw conclusions about the conversion rate. In conclusion, we identified GRA12 as a common virulence factor and validated its importance in clonal and atypical strains.

**Figure 2.**
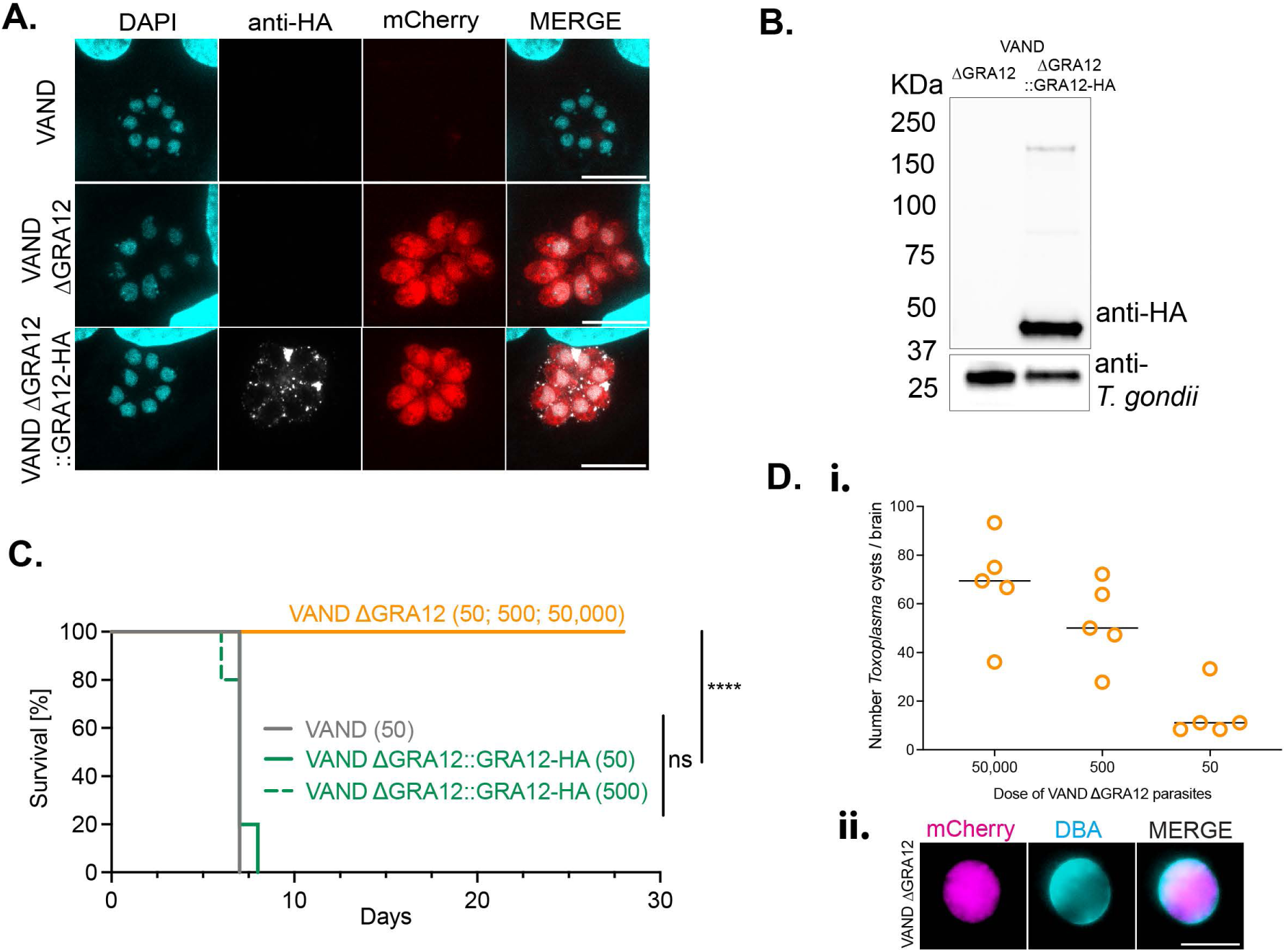
GRA12 is a virulence factor in the highly virulent South American VAND strain *in vivo*. (A) Immunofluorescence localisation of the C-terminal HA-tagged GRA12 in the VAND ΔGRA12::GRA12-HA strain. Scale bar represents 10 µm. (B) Verification of GRA12 expression by anti-HA detection in the VAND ΔGRA12::GRA12-HA strain via western blot. (C) Survival curve of PWD/Phj mice infected with either VAND, ΔGRA12 or ΔGRA12::GRA12-HA strains, dose in parenthesis. Significance was tested using the Mantel-Cox test, N=2-3 mice per group, p **** < 0.0001. (D) Number of brain cysts recovered from PWD/Phj mice infected with VAND ΔGRA12 parasites (i) and immunofluorescence detection of a representative brain cyst stained with DBA. Scale bar represents 50 µm (ii).

### GRA12 is essential to survive the IFNγ-mediated clearance in murine macrophages of different genetic backgrounds

Macrophages are the major cell type infected in the peritoneum during the acute phase^15^. To identify genes specifically important for *Toxoplasma* survival in macrophages, we performed a CRISPR screen using the type I RH KO pool in PWD/Phj mice bone marrow derived macrophages (BMDMs) with or without IFNγ pretreatment (Figure 3A). We used the RH pool to prevent the spontaneous encystation commonly seen for VAND, and PWD/Phj mice to validate parasite factors required in IRG resistant mice. All raw and normalised read counts, L2FC and DISCO scores are reported in Table S7 and the sgRNA dropout rate at each passage in Figure S3A. In line with the *in vivo* screens, GRA12 deletion resulted in the highest fitness defect (Figure 3B). While the strong phenotype observed for GRA12 deletion may have skewed the other effectors phenotypes towards a more neutral score, GRA61, TGGT1_309560, TGGT1_243690, ROP18 and GRA45 also appear to play a role in macrophage survival, but to a lesser extent.

**Figure 3.**
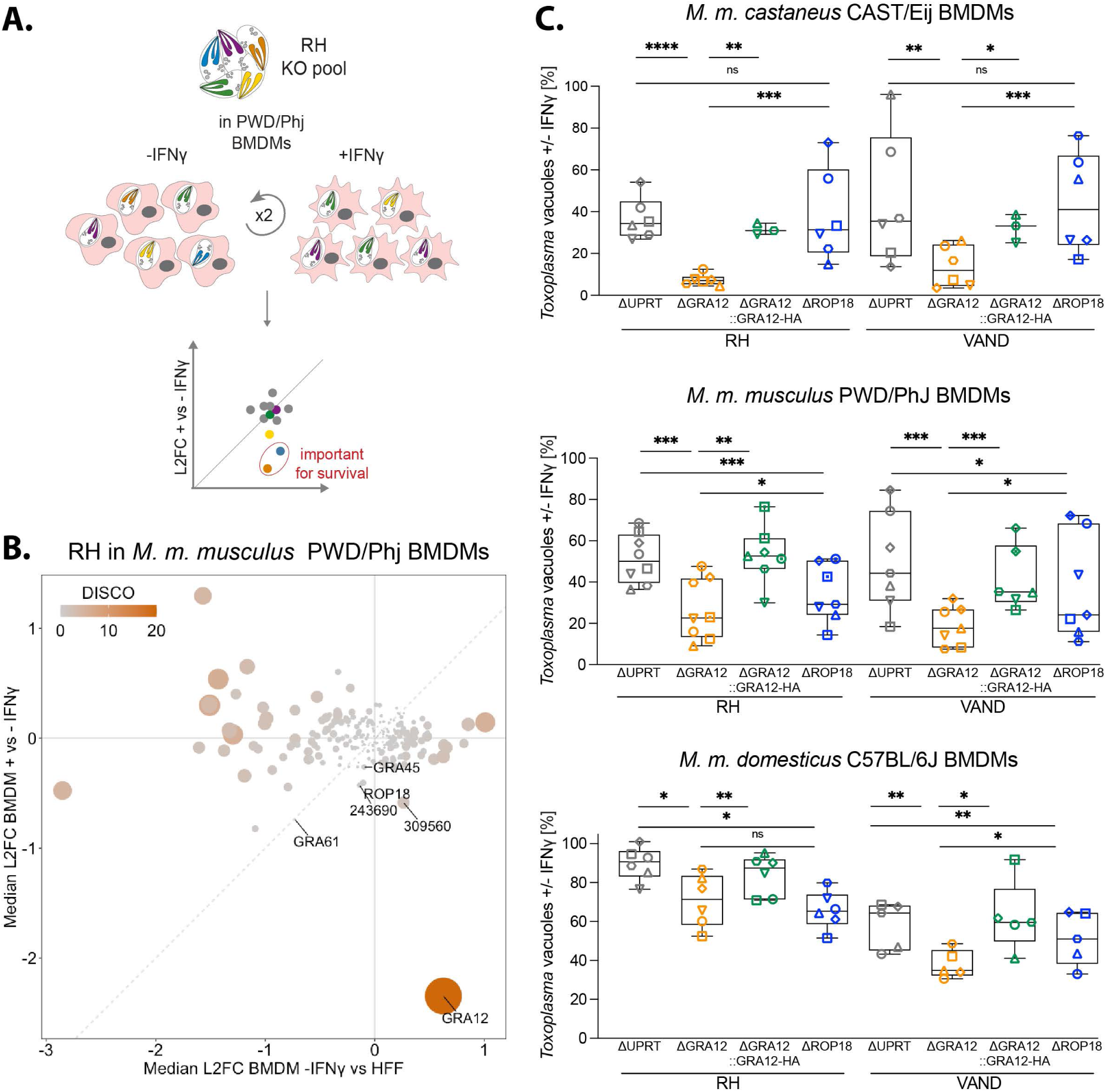
GRA12 is the first identified virulence factor to survive the IFNγ-mediate clearance in different murine subspecies. (A) Scheme of the CRISPR screen of type I RH secretome in PWD/Phj BMDMs to identify *Toxoplasma* factors important to survive IFNγ-mediated restriction. The KO pool was used to infect BMDMs either untreated or pretreated with IFNγ for two consecutive lytic cycles. Surviving parasites were expanded in fibroblasts prior to gDNA extraction and sgRNA amplification for sequencing. The relative abundance of each guide (L2FC) in IFNγ-restricted versus untreated parasites indicates the importance for *in vitro* survival of the relative *Toxoplasma* gene. (B) Scatter plots of the median L2FCs for each gene *in vitro* in CRISPR screens of the RH KO pool in PWD/Phj BMDMs, treated or not with IFNγ. The grey line indicates equal L2FCs. (C) Quantification of high content-automated imaging of parasite vacuoles in IFNγ-treated BMDMs relative to untreated controls. BMDMs of different mouse subspecies were infected with RH or VAND ΔUPRT, ΔGRA12, ΔGRA12::GRA12-HA and ΔROP18 and vacuoles were quantified at 24 h after infection. Symbol shapes indicate biological repeats. Significance was tested using the One-way Anova test with the Benjamini, Krieger and Yekutieli FDR correction. p value * <0.05, ** <0.01, *** <0.001, **** <0.0001.

We next validated the results in macrophage restriction assays using newly generated knockout strains of UPRT (which mimic WT conditions), ROP18 and GRA12, together with the GRA12::HA complemented parasite line (Figure S3B-H). Restriction assays were performed using high-content imaging of BMDMs from *M. m. domesticus* C57BL/6J mice, *M. m. musculus* PWD/Phj mice and *M. m. castaneus* CAST/Eij mice, and parasite burden was quantified on the cumulative mCherry signal of mutant parasites following previously established protocols ^43,44^. At 24 hours post infection (hpi), ΔGRA12 parasites were consistently cleared more than parental strains in IFNγ-primed conditions, which was rescued in the GRA12-complemented line (Figure 3C and S4A). As expected, ROP18 plays no role in conferring protection against the IFNγ response in CAST/Eij BMDMs (Figure 3C and S4A). ROP18 contributed to parasite survival in PWD/Phj BMDMs, albeit to a lesser extent than GRA12. This result matches a ROP18 intermediate phenotype observed in the CRISPR screen in PWD/Phj *in vivo*, but is in contrast with previous work ^39^. Contrary to previously published data, this mouse strain was also susceptible to type I GT1 strain infection (Figure S4B). Sanger sequencing of the PWD/Phj mice confirmed that the IRG genes known to be important for parasite restriction (*Irgb6*, *Irgm1*, *Irgb2-b1*) carry the resistant alleles (data not shown), indicating that an as yet unidentified factor in these mice confers susceptibility to ROP18 expressing parasites. Given that the phenotypic behaviour diverges from published data, results in the PWD/Phj mouse strain warrant further investigation, but is beyond the scope of this study.

To explore GRA12 function across different host species, we performed restriction assays in Wistar rat BMDMs and human foreskin fibroblasts, but no significant change was observed (Figure S4C-D and ^44,52^). As previously reported, GRA12 contributes to IFNγ-mediated parasite survival in murine fibroblasts, although to a lesser extent that in BMDMs (Figure S4D and ^53^). Overall, our data indicate that GRA12 is the first virulence factor validated in different mouse subspecies and across parasite strains, but has probably no function under these conditions in rats or humans.

### Lack of GRA12 results in host cell death by regulated necrosis following Toxoplasma infection

To assess if deletion of GRA12 leads to an increase in host cell death upon IFNγ treatment, we measured the uptake of cell impermeable propidium iodide into the host cell over time. Macrophage cell death was increased in cells infected with ΔGRA12 parasites of either VAND, RH and PRU strains at 9 hpi when compared to controls (15% - 34%) and rescued by the parental and GRA12-complemented strains (Figure 4A-B and S5A-B).

**Figure 4.**
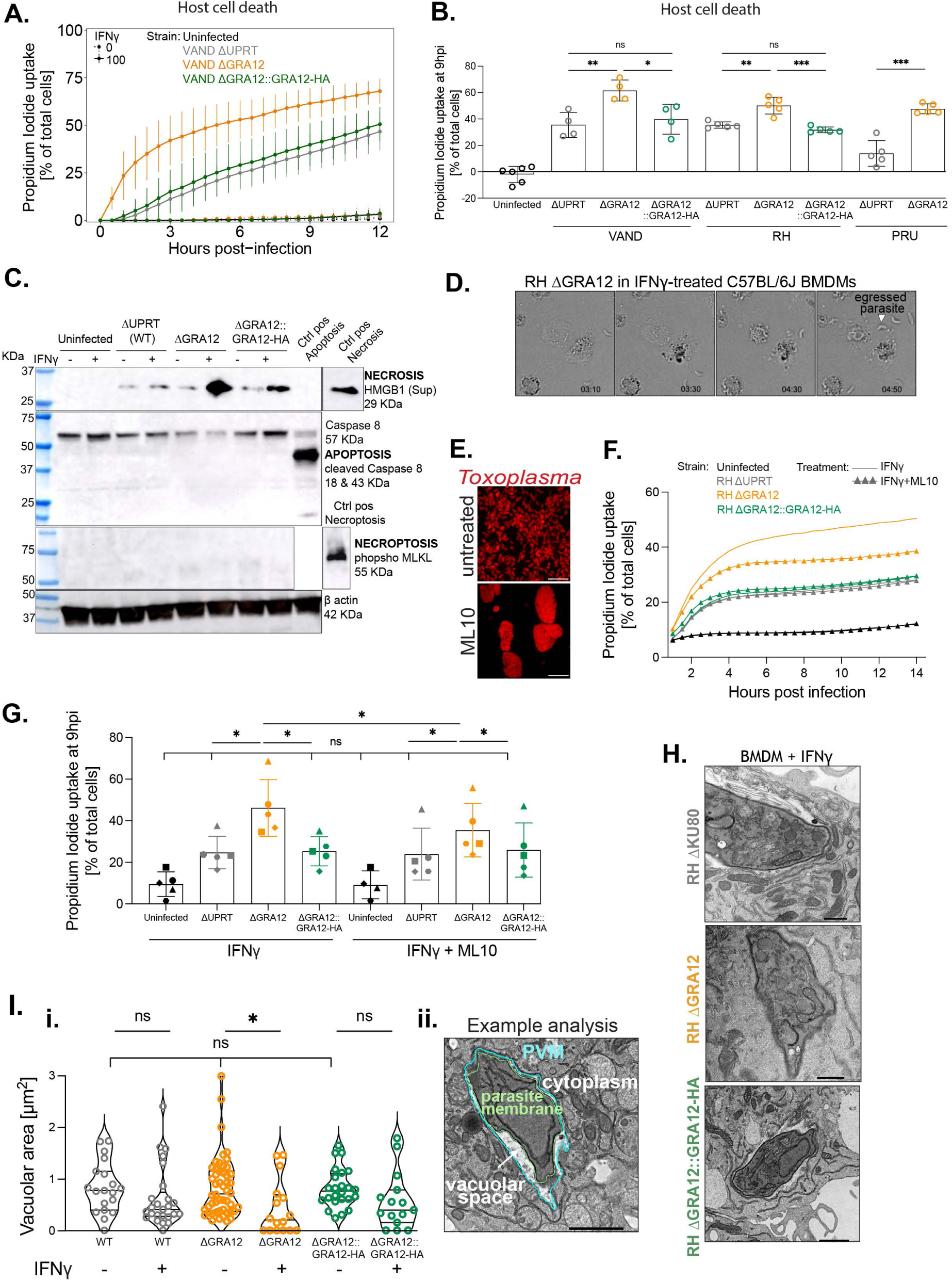
Lack of GRA12 results in IFNγ-mediated host cell death by necrosis and vacuolar collapse. (A) Time course of PWD/Phj BMDMs cell death quantified via Propidium Iodide uptake. Cells were pre-stimulated for 24 h with IFNγ (continuous line) or left untreated as control (dashed line) and then infected with VAND ΔUPRT, ΔGRA12, ΔGRA12::GRA12-HA or left uninfected. The number of dead cells is expressed as a percentage of the total at each time point. (B) Cell death quantification at 9 hpi of IFNγ-treated PWD/Phj BMDMs infected with ΔUPRT, ΔGRA12, ΔGRA12::GRA12-HA in the PRU, RH and VAND strains, or left uninfected as control. Significance was tested using the One-way Anova test with the Benjamini, Krieger and Yekutieli FDR correction. (C) Western blot analysis of the supernatant and cell lysate of C57BL/6J BMDMs pre-treated with IFNγ and infected for 8 h with RH ΔUPRT, ΔGRA12, ΔGRA12::GRA12-HA, or left untreated as control. (D) Live cell imaging of IFNγ-treated C57BL/6J BMDMs infected with RH ΔGRA12 parasites. Time post infection reported as hh:mm in the bottom right corner. Filled arrowheads indicate a parasite egressed from the dying cell. Complete video is in Suppl Movie 1. (E) Fluorescence microscopy detection of RH ΔUPRT parasites after 48 h growth in HFFs either treated or not with 1 µM ML10. Scale bar represents 20 µm. (F) Time course of C57BL/6J BMDM cell death quantified via Propidium Iodide uptake. Cells were pre-stimulated for 24 h with IFNγ, and infected with RH ΔUPRT, ΔGRA12, ΔGRA12::GRA12-HA before 1 hpi treatment with 1 µM ML10 (triangle symbol), or left untreated as control (continuous line). The number of dead cells is expressed as a percentage of the total at each time point. (G) Cell death quantification at 9 hpi of the ML10 treatment experiment. Significance was tested using the One-way Anova test with the Benjamini, Krieger and Yekutieli FDR correction. Symbol shapes indicate biological repeats. (H) Example TEM images of IFNγ-treated or untreated PWD/Phj BMDMs infected with RH ΔKU80, ΔGRA12, ΔGRA12::GRA12-HA for 2 h. (I) Quantification of the vacuolar area from the TEM images. Individual vacuoles were randomly imaged in a single biological experiment. (i) Scatter plot of quantified area. When the PVM was not visible, the vacuolar area was imputed to 0.01 µm^2^. Significance was tested using the Kruskal-Wallis test, (ii) Example of analysis for the vacuolar space quantification on a parasite infecting a BMDM. p * <0.05, ** <0.01, *** <0.001.

Based on the increased cell death in GRA12 deletion mutants, we investigated which cell death pathway is activated upon ΔGRA12 infection. A ∼4-fold increase of the necrosis-associated high mobility group B1 protein marker (HMGB1) was found in the supernatant of ΔGRA12 infected macrophages compared to controls (Figure 4C, quantification in Figure S5C). Cells did not display features of apoptosis or necroptosis, as indicated by lack of cleaved caspase 8 and phosphorylation of MLKL respectively (Figure 4C). Live time-lapse microscopy of IFNγ-treated BMDMs infected with ΔGRA12 parasites showed a sudden shrinkage followed by “burst”-like cell death, with parasites emerging from bursting cells (Figure 4D and Suppl Movie 1). Numerous extracellular parasites were observed specifically in IFNγ-treated macrophages infected with ΔGRA12 parasites (Figure S5D). Therefore, we hypothesised that the lack of GRA12 might promote host cell necrosis caused by early egress of the parasite. Consequently, we treated parasites with a cGMP-dependent protein kinase (PKG) inhibitor developed for *Plasmodium falciparum* parasites, ML10 ^54^. ML10 treatment blocked parasite egress (Figure 4E), indicating that the compound also functions in *Toxoplasma.* ML10 treatment reduced parasite numbers, but did not interfere with IFNγ-mediated restriction (Figure S5E). ML10 treatment partially reduced the propidium iodide uptake in ΔGRA12 infected macrophages in a IFNγ-dependent manner (Figure 4F-G and S5F), suggesting that early egress of ΔGRA12 parasites contributes to increased necrotic cell death, but is not the sole driver.

Transmission Electron Microscopy (TEM) of infected BMDMs revealed that ΔGRA12 parasites have significantly reduced vacuolar space in IFNγ-treated macrophages compared to the untreated condition. In over 30% of cases the PVM is not clearly separated from the parasite plasma membrane (Figure 4H-I). Vacuole collapse has been associated with nitric oxide (NO) production and GBP2 recruitment ^32^. However, infection with ΔGRA12 parasites neither increases NO levels (Figure S5G) nor is restriction rescued by the iNOS inhibitor 1400W (Figure S5H-I). We also observed no increase in GBP2 recruitment to the PV in ΔGRA12 infected cells, or other host proteins previously shown to restrict *Toxoplasma* (IRGd, IRGb10, ubiquitin and p62 in Figure S6, similarly to published data on IRGa6 and IRGb6 ^53,55^) ^56^. GRA12 function depends on IRGM1 and IRGM3 ^53^, however their cellular distribution appears similar across strains (not shown). In summary, these data suggest that lack of GRA12 increases host necrotic death via a yet undescribed mechanism.

### GRA12 homologues from closely related Coccidian parasites rescue *Toxoplasma* ΔGRA12 restriction

GRA12 is predicted to contain a signal peptide, a short N-terminus, a putative transmembrane domain, and a larger 36 KDa C-terminal domain ^57^. To test if the N- or C-terminus may face the host cell cytoplasm where they could directly interact, and interfere, with host cell proteins, we assessed the topology of GRA12. We generated an endogenously C-terminally HA-tagged line (RH GRA12-HA) and a complemented N-terminally HA-tagged line (RH ΔGRA12::HA-GRA12, Figure S7A-E). We used low concentrations of the mild detergent saponin to permeabilise only a fraction of PVMs, and localise the GRA12 termini in either the cytoplasmic or vacuolar compartments via immunofluorescence detection with an anti-HA antibody. HA signal was only detected in vacuoles that were also anti-*Toxoplasma* antibody positive, indicating that both ends of GRA12 face the luminal side of the PV and therefore do not face the host cell cytosol (Figure 5A). As previously reported, GRA12 predominantly co-localises with the IVN protein GRA2 and only partially with the PVM marker GRA3, further supporting a model in which GRA12 is not directly interacting with host proteins on the PVM surface (Figure S7F) ^53,57^.

**Figure 5.**
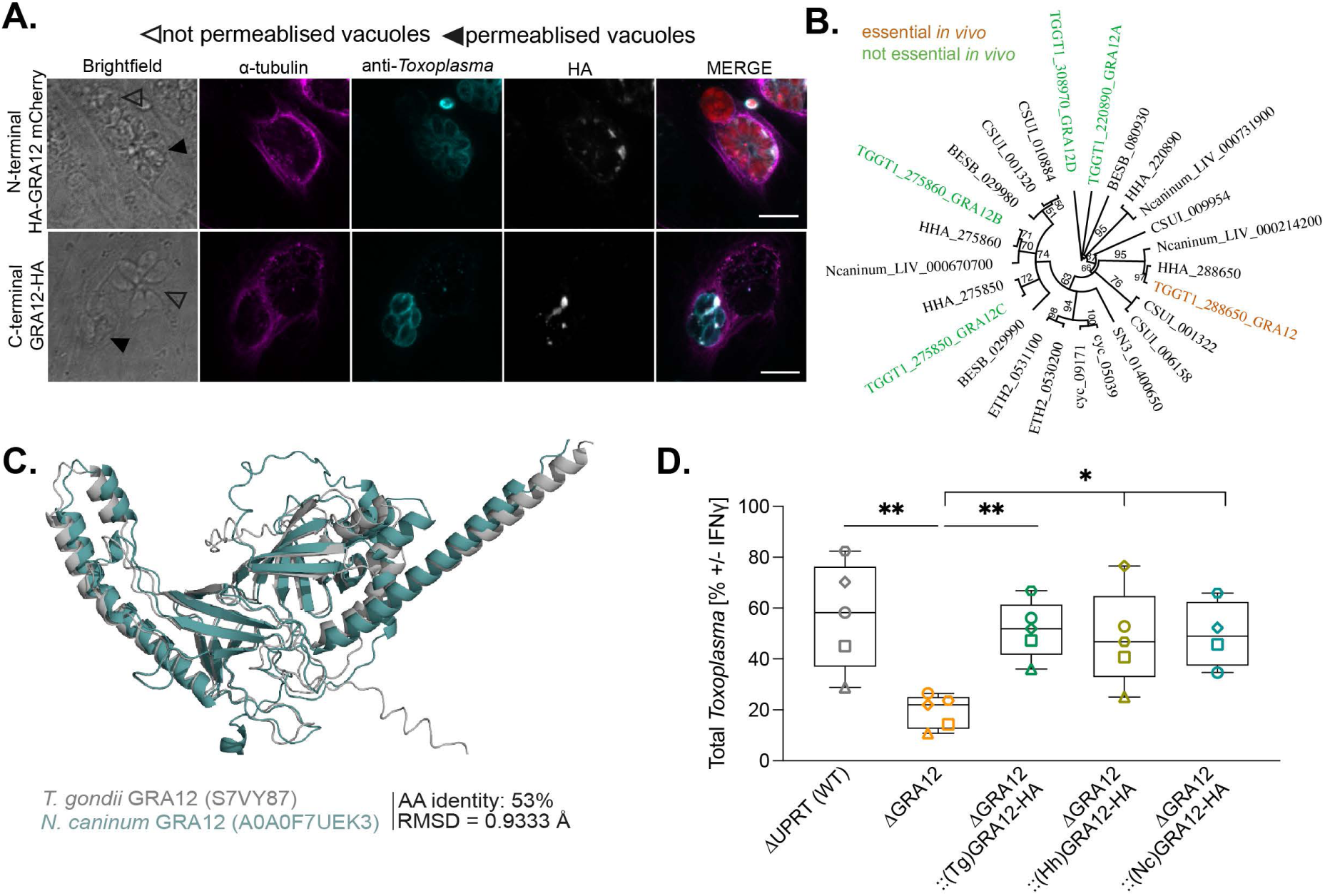
GRA12 is an intravacuolar protein conserved across Coccidia. (A) Immunofluorescence localisation of GRA12 in partially permeabilised human fibroblasts infected with the N-terminally tagged line (upper panel) or the C-terminally tagged line (lower panel). Filled arrowheads indicate a permeabilised vacuole and empty arrowheads indicate a non-permeabilised vacuole. Scale bar represents 10 µm. (B) Neighbor-Joining tree of Coccidian parasites GRA12 homologues built with Bootstrap method and considering GRA12D as outgroup. The bootstrap support values, based on 100 replicates, are indicated above each node. (C) Structural overlay of *Toxoplasma* GRA12 (grey) and *N. caninum* GRA12 homologue (teal) as predicted by AlphaFold2. Root Main Square Deviation (RMSD) was calculated using PyMol. (D) Relative *Toxoplasma* growth in IFNγ-treated versus untreated BMDMs. BMDMs were infected with the RH ΔGRA12 strain, or the RHΔGRA12 strain complemented with *Toxoplasma* (Tg) GRA12 or the *H. hammondi* (Hh) or *N. caninum* (Nc) GRA12 homologues, or the RH ΔUPRT strain as control for 24h before a plate reader quantification of the mCherry signal as proxy for parasite growth. Significance was tested using the One-way Anova test with the Benjamini, Krieger and Yekutieli FDR correction, p* < 0.05, ** < 0.01.

To identify putative GRA12 interaction partners, we performed protein co-immunoprecipitation (co-IP) in IFNγ-treated murine macrophages infected with GRA12-HA or wildtype parasite as control and analysed the samples by mass spectrometry. As expected, GRA12 showed the highest enrichment (Figure S7G-H and Table S8), but only a few putative host interactors were identified: the murine ATPase plasma membrane Ca2+ transporting 1 (ATP2B1) and the mitochondrial protein Prohibitin 2 (PHB2). However, control IPs and immunofluorescence assays failed to confirm their specific enrichment at the vacuole or reciprocal co-immunoprecipitation with GRA12 (data not shown). Previously identified interactors of GRA12, such as GRA2, ROP5 and ROP18 ^57,58^ were not identified in this experiment. These may have been missed here because of technical differences between pull down protocols. But given that GRA12 is important even in strains where ROP5 and ROP18 have no protective role, these interactions are probably not important for GRA12 function.

GRA12 has four paralogues (GRA12A-D) which share an overall pairwise amino acid sequence identity below 30% and that are, except for GRA12C, highly expressed in both tachyzoite and bradyzoite stages. All Coccidia parasites have 1 to 4 *Gra12*-like orthologues (Figure 5B). We and others have shown that only GRA12, but not its paralogues, is essential in *Toxoplasma* infections *in vivo* ^49–51^.

The *Hammondia hammondi* and *Neospora caninum* orthologues closest to *Toxoplasma* GRA12 share 85% and 53% of the amino acid sequence respectively. The GRA12 protein structure prediction by Alphafold suggests a conserved fold, even with the more diverse *Neospora* homologue (Figure 5C). This suggests a similar function of GRA12 in these related species. To test this hypothesis, we complemented RH ΔGRA12 with HA-tagged orthologues of *Hammondia hammondi* (HHA_288650) or *Neospora caninum* (Ncaninum_LIV_000214200, Figure S7I-J). Both proteins localised within the vacuole, similar to the *Toxoplasma* GRA12 (TgGRA12, Figure S7K) and rescued IFNγ-mediated restriction of RH ΔGRA12 parasites in restriction assays (Figure 5D). This suggests that GRA12 also functions to prevent the IFNγ-mediated clearance in closely related Coccidian parasites.

## Discussion

*Toxoplasma* can infect and persist in a wide range of hosts. While past research has focused on secreted virulence factors that render some strains more pathogenic than others, no protein that allows the parasite to persist in all hosts has been identified yet. Here, using systematic *in vivo* CRISPR-Cas9 screens, we identified secreted virulence factors that function across *Toxoplasma* strains and mouse genotypes. Among these, deletion of GRA12 consistently had the most pronounced effect on parasite survival *in vivo*.

We show that the absence of GRA12 leads to a rapid death of the infected cell. Early egress partially explains the increase in host cell death, and time-lapse imaging suggests that it is probably not preceding it. Parasite egress is more likely a response to the rapid induction of the host immune response, causing the activation of regulated cell death pathways, cellular shrinkage and changes in intracellular ion concentrations ^59^, triggering egress. The observed host cell burst does not resemble cell death mechanisms previously associated to *Toxoplasma* infection, such as pyroptosis and necroptosis, which are characterised by an initial swelling phase ^60^. This indicates that probably a different cell death pathway is induced. We did not observe or test the ability of egressed parasites to successfully infect neighbouring cells. However, the strong reduction of the ΔGRA12 parasites both *in vivo* and *in vitro* suggests that the combined effect of host niche destruction and enhanced parasite egress is sufficient to halt the infection. Since GRA12 does not seem to span the membrane to interact with host proteins, and we did not identify convincing protein interaction partners, we hypothesise that GRA12 might have a role in stabilising the PVM. As a result, the deletion of GRA12 might establish hypersensitivity to clearance mechanisms, such as vacuolar breakage, increasing egress and triggering of host-cell death pathways. The collapsed vacuolar space of ΔGRA12 parasites would support this hypothesis.

Strikingly, GRA12 orthologues from closely related parasites rescued GRA12 function, indicating that its function might be conserved in these other Apicomplexa. It would be interesting to assess how far the conservation of GRA12 extends by testing whether the most distant GRA12 homologue, which is in Eimeria *spp.* (25% shared amino acid sequence), is still able to complement the *Toxoplasma* GRA12. Type II interferon is also important for clearance of Coccidian parasites, including *N. caninum* and *H. hammondi* ^61,62^, supporting a similar function of GRA12 to survive the IFNγ-mediated clearance of both parasite species. However, while the murine host is key in the life cycle of *H. hammondi* ^62^, *N. caninum* only sporadically infects mice in nature ^63^, suggesting that the function of GRA12 might not be exclusive to murine cells.

In addition to GRA12, we found that deletion of GRA45, GRA61, GRA50 and GRA23 also reduces parasite survival *in vivo* in all conditions tested here. GRA45 has previously been shown to act as a chaperone to insert proteins into the PVM, for example GRA23 which explains the presence of both proteins as hits ^47^. While GRA12 was found to have a partial membrane association with the PV membrane ^57,64^, for which GRA45 may be required, our data shows that both the N- and C-termini of GRA12 localise within the PV. This suggests that GRA12 does not span the membrane and probably functions independently of GRA45. The function of GRA50 and GRA61 is unknown, and future studies should explore a possible function of these proteins. However, our proteomics data did not identify them as interaction partners of GRA12, suggesting that they probably function independently of GRA12, or at least not in a complex. Although we may have missed some exported proteins in our study for technical reasons (i.e. not enough datapoints or missing annotation as exported protein at the time of the study ^65^), we likely identified the vast majority of virulent factors important in the tested conditions. Our results, together with previous studies that focussed on virulence factors in human cells where GRA12 does not play an important role ^44,52,66^, suggest that individual virulence factors have evolved to protect the parasite in different species.

We also identified strain-specific effector proteins. Most of these have not been discussed here, but ROP18 behaves as expected, lending support to the robustness of the screens: it is required to confer high virulence to RH and VAND strains in susceptible murine subspecies, is less important for PRU survival and is dispensable in VEG. Surprisingly, the three recently identified secreted members of a protein complex required to prevent parasite IFNγ-mediated clearance in human fibroblasts (GRA57, GRA70, GRA71) ^44,52^ show a fitness defect in the VAND strain isolated from a lethal human infection, but not in any other strain. This is an important result and warrants further investigation. Positive selection of this protein complex in a zoonotic reservoir could explain the increased virulence of South American *Toxoplasma* strains in humans.

## Material and methods

### Animal work

All mouse work was approved by the UK Home Office (project licence P1A20E3F9) and the Francis Crick Institute Ethical Review Panel and carried out in accordance with the UK Animals (Scientific Procedure) Act 1986 and European Union directive 2010/63/EU. Different strains of *Mus musculus* inbred laboratory mice were used: the C57BL/6J strain, the *M. m. musculus* PWD/Phj strain and the *M. m. castaneus* CAST/Eij strain. Non-regulated procedures were performed on Wistar rats (kindly provided by the Biology Research Facility of the Francis Crick Institute) for tissue collection. All animals were bred and housed under pathogen-free conditions in the biological research facility at the Francis Crick Institute.

Bone Marrow Derived Macrophages (BMDMs) were derived from all murine inbred laboratory strains mentioned above as well as from Wistar rats. Animals were sacrificed by cervical dislocation and the bone marrow collected for extraction of monocytes with a protocol provided by the Wack lab. Bone marrow cells were seeded in 15 cm diameter bacterial Petri dishes (Falcon) and differentiated into BMDMs for 6-7 days with complete RPMI 1640 Medium ATCC modification (Gibco A1049101) supplemented with 10% heat-inactivated FBS (Gibo 10500064), 100 U/ml Penicillin-Streptomycin (Gibco), 50 μM 2-mercaptoethanol (Gibco) and 20% L929 conditioned media. Following differentiation, BMDMs were resuspended in a solution of cold PBS with 2% FCS and 2 mM EDTA (ThermoFisher), and either used for experiments with complete medium without 2-mercaptoethanol or frozen in 9:1 FCS/DMSO until use.

### Cell culture and parasite strains

Primary HFFs and MEFs (ATCC) were maintained in Dulbecco’s modified Eagle’s high glucose and GlutaMAX^TM^ supplemented medium (DMEM 61965059) with 10% FBS at 37 °C and 5% CO2. *Toxoplasma* strains VAND (gift from Eva Frickel), VEG (gift from Martin Blume), RH ΔHXGPRT ^67^, PRU ΔHXGPRT (gift from Dominique Soldati, as in ^67^) and RH ΔKU80 ^68^ were maintained by growth in confluent HFFs and passaged every 2–3 days. Parasites were regularly tested for *Mycoplasma* spp. contamination at the in-house Cell Services Facility. To prepare extracellular parasites for infection, infected HFFs were syringe lysed with a 23G needle, filtered through a 5 μm sterile filter (Millipore) to remove debris and counted in a haemocytometer chamber prior to dilution to the desired concentration.

### CRISPR-Cas9 screens *in vivo* and *in vitro*

CRISPR-Cas9 screens were performed according to previously optimised protocols ^42–44^ and described as follow. All primers used in this work are listed in Table S1.

### Library generation

Screens in RH ΔHXGPRT and PRU ΔHXGPRT strains were performed with a sgRNA library cloned into the previously created vector pCas9-GFP-HXGPRT::sgRNA ^69^. For screens in the VAND and VEG strains a novel vector pCas9-GFP-DHFR-TS::sgRNA was created by Gibson cloning the DHFR-TS selection cassette amplified with primers 1-2 from the template GRAx-TGGT1-069070-DHFR-TS in pCas9-GFP-HXGPRT::sgRNA double-digested with SmaI/EcoRI (NEB). PCRs were performed with the proof-reading polymerase KOD (Sigma-Aldrich), and plasmids verified by sequencing with primers 3-7. The pool of ssDNA oligonucleotides encoding the protospacer sequences was selected from an arrayed library using an Echo 550 Acoustic Liquid Handler (Labcyte) in three independent events to minimise imprecisions in the automated liquid handling, and then pooled. The pooled oligonucleotides were integrated in both CRISPR vectors by Gibson cloning after digestion with PacI/NcoI (NEB), resulting in pool libraries of 1240 sgRNAs and 1247 sgRNAs for the HXGPRT and DHFR-TS vectors respectively, and an average of 5 sgRNAs/gene. Individual libraries were sequenced by Illumina sequencing as previously described ^42^.

### CRISPR pool creation

Parasite transfection was performed in 3-5 replicates and each replicate was sequenced to ensure transfection reproducibility. Libraries were linearised with KpnI-HF or NheI-HF (NEB) for the HXGPRT or DHFR-TS vectors respectively, with an overnight digestion prior to phenol-chloroform purification and transfection with the P3 Primary Cell 4D-Nucleofector kit (Lonza V4XP-3032) in a Amaxa 4D Nucleofector (Lonza AAF-1003X) with the program EO-115. Integration of the pCas9-GFP-HXGPRT::sgRNA or pCas9-GFP-DHFR-TS::sgRNA libraries was induced upon treatment with 25 µg/ml Mycophenolic acid (Sigma-Aldrich) and 50 µg/ml Xanthine (Sigma-Aldrich) the following day, or with 2 µM Pyrimethamine (PYR, Merck) 2 h post transfection respectively. Three days post transfection, parasites were syringe lysed and added to fresh HFFs monolayers with 100 U/ml Benzonase (Merck) overnight to remove traces of input DNA. Six to eight days post transfection, samples were taken for gDNA preparation or used as *in vivo*/*in vitro* inoculum.

### *In vivo* selection

Different combinations of parasite and mouse strains were used as specified throughout the manuscript. The inoculum was diluted at 1E6 parasites/ml in PBS and mice were injected intraperitoneally with 200 μl equal to 2E5 parasites for all but VEG, for which a dose of 5E5 parasites was used to account for higher restriction of this parasite strain *in vivo*. Eight-to sixteen-week-old male mice were injected, and each *in vivo* screen was performed on five mice. Five days post infection, parasites and peritoneal cells were isolated by double peritoneal lavage with 5 ml of PBS. Peritoneal cells were pelleted at 500 *g* for 5 min, resuspended in PBS and intracellular parasites were syringe lysed with 27G and 30G needles and used to infect HFFs in DMEM supplemented with 100 U/ml Pen/Strep. After one lytic cycle, expanded parasites were pelleted for gDNA extraction.

### *In vitro* selection

BMDMs from PWD/Phj mice were used for the RH ΔHXGPRT screen to match the mouse strain-parasite strain combination used in the *in vivo* screen. 7E6 BMDMs were seeded and treated for 24 h with 10 U/ml murine IFNγ (Gibco PMC4031) or left untreated prior to infection. To limit co-infections, 1.4E6 parasites per dish corresponding to a MOI 0.2 and a coverage of 1,230 parasites/sgRNA were used. At 48 hpi BMDMs were scraped, syringe lysed with a combination of 23G, 27G and 30G needles, and parasites from round 1 were used at a MOI 0.2-0.3 for infection of round 2 in BMDMs similarly prepared. Each condition was performed in triplicate. Similarly to the *in vivo* screen, parasites were expanded for one lytic cycle in fibroblasts, before gDNA extraction and sgRNA amplification for sequencing.

### Illumina sequencing of sgRNA and data analysis

Genomic DNA was extracted from samples using DNEasy Blood kit (Qiagen), then guide sequences were amplified by nested PCR using KAPA HIFI Hotstart PCR kit (Kapa Biosystems KK2501) and KAPA Pure beads (Kapa Biosystems KK8000). Primers 8-27 were used for nested PCRs. Purified PCR products were then sequenced on a HiSeq400 (Illumina) with single end 100 bp reads at a minimum read depth of 5 million reads/sample. gRNA sequences were aligned to the reference library and counts were normalised using the median of ratios, then sgRNA not present in all samples or in less than three peritoneal samples were removed from the analysis. For the *in vivo* CRISPR screens, the median L2FC for each sgRNA was calculated from the normalised counts in the inoculum sample relatively to the plasmid (Median L2FC *in vitro*), or from the peritoneal samples at day five compared to the inoculum (Median L2FC *in vivo*). For the *in vitro* CRISPR screens, the median L2FC for each sgRNA was calculated from the normalised counts in the unstimulated BMDMs relative to the inoculum (Median L2FC BMDM-IFNγ vs HFF), or from the IFNγ-stimulated BMDMs relative to unstimulated BMDMs after the second round of selection (Median L2FC BMDM + vs - IFNγ). The median absolute deviation (MAD) score across gRNA L2FCs targeting each gene was calculated, and genes with the highest 1.5% of MAD scores were removed from the analysis. A DISCO score based on the local FDR-corrected q-value was calculated for each L2FC.

### Creation and validation of the VAND mutant strains

To create VAND mutant strains, the ProGRA1::mCherry::T2A::HXGPRT::TerGRA2 plasmid ^42^ encoding the repair template mCherry::T2A::HXGPRT was modified by replacing *Hxgprt* with the Chloramphenicol acetyltransferase (CAT) sequence from the pG140::DiCre plasmid ^70^, amplified with primers 28-31 and Gibson cloned. The resulting ProGRA1::mCherry::T2A::CAT::TerGRA2 plasmid was validated with primers 3 and 32. VAND ΔGRA12, ΔROP18 and ΔUPRT strains were created by co-transfecting VAND with the protospacer-encoding pCas9::GRA12, pCas9::ROP18 and pCas9::UPRT plasmids (created with primers 33-35 or with the original pCas9::UPRT plasmid, and validated by sequencing with primer 36) and the relative repair templates amplified with primers 37-42 from the ProGRA1::mCherry::T2A::CAT::TerGRA2 plasmid. The disruption of the endogenous locus was PCR validated with primers 43-48 and by Nanopore sequencing as follow. Three libraries were prepared from 200 ng of genomic DNA per sample using the Nanopore SQK-RBK114 kit. Sequencing was carried out on a PromethION flow cell (R.10.4.1) using a P2 Solo device for 24 h. Basecalling was performed with Dorado v0.7.3 (<u>Dorado</u>), utilising the “sup” model, and the reads were aligned to the ToxoDB-65_TgondiVAND genome assembly reference from ToxoDB release 49 (<u>VAND</u>). The aligned reads were indexed and sorted for future visualisation. Post-demultiplexing, Dorado was used again to detect and trim primers and adapters from the reads. Mapped reads were visualised against the reference genome using IGV v2.16.2 ^71^. Nanopore sequence reads alignment showing disruption of the endogenous loci in VAND ΔGRA12, ΔROP18 and ΔUPRT strains, and integration of the repair template is represented in Figure S8.

To create the complemented strain VAND ΔGRA12::GRA12-HA, the 5’UTR and cDNA of GRA12 (TGVAND_288650) were Gibson cloned in the pUPRT vector ^43^ with primers 49-54, and the plasmid pUPRT::GRA12 was verified by sequencing with primers 41, 49-50. VAND ΔGRA12 parasites were co-transfected with the ScaI (NEB) digested pUPRT::GRA12 plasmid and the pCas9::UPRT plasmid, and selected for integration in the *Uprt* locus with 5 μM 5′-fluo-2′-deoxyuridine (FUDR, Sigma F0503) the day after transfection. A clonal population of the resulting VAND ΔGRA12::GRA12-HA parasites was validated by PCR with primers 57-60, by western blot and immunofluorescence detection of the protein. PCRs for cloning were performed with the proof-reading polymerase ClonAmp HiFi PCR Premix (Takara) and PCR for validation of strains were performed with EmeraldAmp GT Master Mix (Takara).

### In vivo infections

PWD/Phj mice 8-12 weeks old were infected intraperitoneally with different doses of parasites in 200 µl PBS. Immediately after infection, fitness of the parasites used as inoculum was assessed via plaque assay to establish the precise parasite dose. Mice were monitored and weighed regularly for the duration of the experiments or until reach of the humane endpoint. To determine the number of cysts in the brain of infected animals, mice were euthanised at 28 days post infection. The brain was homogenised in 1 ml PBS and stained with Fluorescein-conjugated Dolichos Biflorus agglutinin (1:200, Vector Laboratories RL-1031) for 1 h at room temperature. Fluorescently labelled cysts were counted using a Ti-E Nikon microscope with a 40x magnification in a 1:1 dilution of homogenate and PBS. At least one third of the brain homogenate was analysed for cysts presence.

### Plaque assays

HFFs were grown to confluency in T25 flasks and infected with 100-200 parasites to grow undisturbed for 10 days for RH derived strains and 14 days for other strains. Up to 25,000 parasites were plaqued to determine the transfection efficiency of the CRISPR transfections. Cells were fixed and stained in a solution with 0.5% w/v crystal violet (Sigma), 0.9% w/v ammonium oxalate (Sigma), 20% v/v methanol in distilled water then washed with tap water. Plaques were imaged on a ChemiDoc imaging system (BioRad) and measured in Fiji ^72^.

### Creation of the RH ΔKU80-derived strains

To create the RH ΔKU80 GRA12 strain, RH ΔKU80 parasites were co-transfected with the pCas9::GRA12 plasmid and the repair template amplified from the ProGRA1::mCherry::T2A::HXGPRT::TerGRA2 plasmid with primers 37-38, and selected with M/X 24 h after transfection. A clonal population of RH ΔGRA12 strain was validated by PCR with primers 43-44, 61-62. The GRA12 C-terminal endogenously tagged RH GRA12-HA strain was created by co-transfection of RH ΔKU80 parasites with the pCas9::GRA12-CT plasmid encoding a protospacer targeting the 3’ UTR (created with primers 33 and 63, and validated with primer 36) and the repair template amplified with primers 64-65 from the HA-TerGRA2::ProDHFR-HXGPRT-TerDHFR plasmid ^43^, and selected with M/X 24 h after transfection. A clonal population of the resulting RH GRA12-HA strain was validated by PCR with primers 57-58, 60 and 66, by western blot and by immunofluorescence detection of the protein.

The GRA12 N-terminally tagged RH ΔGRA12::HA-GRA12 strain and the complemented RH ΔGRA12::GRA12-HA strains with either the *Toxoplasma* (RH ΔGRA12::(Tg)GRA12-HA), *Hammondia hammondi* (RH ΔGRA12::(Hh)GRA12-HA) or *Neospora caninum* (RH ΔGRA12::(Nc)GRA12-HA) homologues were all created by cloning into the *Uprt* locus via co-transfection of pCas9::UPRT and a pUPRT repair plasmid. The plasmid pUPRT::GRA12(GT1) to complement the type I TGGT1_288650 HA-tagged isoform of GRA12 in RH ΔGRA12 and create the RH ΔGRA12::(Tg)GRA12-HA strain, or simply called RH ΔGRA12::GRA12-HA, was created similarly to pUPRT::GRA12 with plasmids 52-54 and 67. In the pUPRT::HA-GRA12 plasmid for N terminal GRA12 tagging, a HA-tag followed by a Gly-Gly linker were inserted at position 47 of the amino acid sequence, between the signal peptide sequence (AA 1-38) and the putative transmembrane domain (AA 93-115) as predicted by ^57^. The plasmid pUPRT::HA-GRA12 was created similarly to pUPRT::GRA12 with primers 50, 67-71. The *H. hammondi* and *N. caninum* cDNA isoforms were PCR amplified from a codon optimised gBlock (IDT, 72 in Table S1) and Gibson cloned to create plasmids pUPRT::(Hh)GRA12 and pUPRT::(Nc)GRA12 with primers 73-80. All complemented plasmid constructs were validated by sequencing with primers 55-56, ScaI-linearised and co-transfected with pCas9::UPRT, and parasites selected for integration by FUDR treatment the day after. All complemented strains were validated by PCR with primers 55-56 and by immunofluorescence detection of the protein. PCRs for cloning were performed with the proof-reading polymerase ClonAmp HiFi PCR Premix (Takara) and PCR for validation of strains were performed with EmeraldAmp GT Master Mix (Takara).

### Immunofluorescence Assay and Live-Cell Imaging

HFFs were seeded on 8-well μ-slides (Ibidi 80806) and infected for 24 h, and BMDMs were seeded on 96-well plates (Revvity PhenoPlate 6005430) and infected for 90 min, before fixation in 4% paraformaldehyde (PFA) for 15 min. Samples were then permeabilised in 0.1% saponin for 10 min and blocked with 3% bovine serum albumin (BSA) in PBS for 1 h. For the localisation of the N and C-terminal extremities of GRA12 in Figure 5A, samples have been treated with 0.001% saponin for 15 s to permeabilise only some but not all PVMs, or fully permeabilised for 10 min or left untreated as controls. All antibodies were incubated in 3% BSA in PBS for 2 h at RT or at 4°C o/n, followed by 3x washes in PBS and a secondary staining combined with DAPI. Images were acquired on a VisiTech instant SIM (VT-iSIM) microscope using a 150x oil-immersion objective with 1.5 μm z axis steps (Figure 2A, Figure S3E and Figure S7F,K), on the 3i Marianas II LightSheet microscope using a 40x or 60x objectives with 8 μm z axis steps (Figure S6 and Figure 5A) or on the Nikon Ti-E inverted widefield microscope with a Nikon CFI APO TIRF 100x/1.49 objective and Hamamatsu C11440 ORCA Flash 4.0 camera running NIS Elements (Nikon) (Figure S2B). Resultant stacks were deconvoluted and processed using Microvolution plugin in Fiji. Primary antibodies used were 1:500 rat anti-HA (Roche 11867423001), 1:1,000 rabbit anti-*T*oxoplasma (Abcam ab138698), 1:1,000 mouse anti-α tubulin (Abcam ab7291), 1:1,000 mouse anti-GRA2 and 1:1,000 rabbit anti-GRA3 (both gifts from Jean François Dubremetz), 1:1,000 rat anti-GRA12 (gift from Maryse Lebrun), 1:1,000 rabbit anti-IRGd6 (081/3) and 1:6,000 anti-IRGb10 (940/6, both gifts from Jonathan Howard), 1:500 rabbit anti-GBP2 (Proteintech 11854-1-AP), 1:200 mouse anti-Ubiquitin (FK2, Merck ST1200), 1:200 rabbit anti-p62 (Genetex GTX111393), 1:100 mouse anti-IRGm1 (Sigma SAB1305514) and 1:100 mouse anti-IRGm3 (Santa Cruz sc-136317). All secondary antibodies used were from Thermo Fisher and were 1:500 anti-rat AlexaFluor 647 (A21247), 1:500 anti-mouse AlexaFluor 647 (A21235), 1:500 anti-rabbit AlexaFluor 647 (A21244), 1:2,000 anti-rabbit AlexaFluor 488 (A11008), 1:2,000 anti-mouse AlexaFluor 488 (A11029), 1:2,000 anti-rat AlexaFluor 488 (A11006), 1:2,000 anti-rabbit AlexaFluor 405 (A-31556).

For the Live Cell Imaging in Figure 4D and Video S1, C57BL/6J BMDMs were seeded in 8-well μ-slides (Ibidi 80806), pretreated for 24h with 100 U/ml mIFNγ or left untreated, and infected with RH ΔUPRT, ΔGRA12 or ΔGRA12::GRA12-HA at a MOI of 1. After 30 min, cells were washed twice and imaged for 12 h with 5 min interval on a 3i Marianas II LightSheet microscope using a 40x objective.

### Western blot

Syringe-lysed parasites from HFFs culture were used for western blots for strain validation, while 1E6 C57BL/6J BMDMs were seeded in 6-well plates to identify cell death pathways by western blots. BMDMs were treated for 24 h with 100 U/ml mIFNγ or left untreated as control, and infected for 8 h with RH ΔUPRT, ΔGRA12, ΔGRA12::GRA12-HA at a MOI 3. BMDMs were treated with 0.25 mM H_2_O_2_ for 4 h, or with 0.5 μM Staurosporine (Cayman 81590) for 8 h or with 20 μM Z-VAD (ApexBio A1902) for 30 min before 7 h treatment with 20 ng/ml TNF-α (Biolegend 718004) and 100 nM SM-164 (Cayman 28632), as positive control for death by necrosis, apoptosis and necroptosis respectively. Proteins from the supernatant were precipitated with 4x ice-cold acetone for 1 h in −20°C, then were spun down at 13,000 *g* for 10 min and the pellet let dry and resuspended in 4x sample buffer solution (Bioworld BW-21420018-3) supplemented with 2-mercaptoethanol. BMDMs were washed once with cold PBS and lysed on ice in RIPA buffer (Thermo Fisher 89901) supplemented with 2× cOmplete Mini EDTA-free Protease Inhibitor Cocktail (Roche) for 30 min, then spun at 13,000 *g* 5 min to remove the insoluble fraction. Samples were added with 4x sample buffer solution supplemented with 2-mercaptoethanol and boiled for 5 min at 95°C together with the supernatants. Samples were separated on a SDS-PAGE on precast 4-15% TGX stain-free gels (Bio-Rad) and the Precision Plus Protein All Blue ladder (Bio-Rad) was used. Proteins were transferred to a nitrocellulose membrane for 7 min using the Turbo midi protocol on the Trans-Blot Turbo transfer system (Bio-Rad). Membranes were blocked in blocking solution (5% milk in PBS with 0.1% Tween-20) for 1 h RT or o/n at 4°C, followed by incubation with primary antibodies in blocking solution either for 2 h RT or o/n at 4°C, and of secondary antibodies for 1 h RT. Primary antibodies used were 1:1,000 rabbit anti-*T*oxoplasma (Abcam ab138698), 1:10,000 rabbit anti-ROP18 (gift from David Sibley), 1:1,000 rat anti-HA Horseradish Peroxidase (HRP) conjugated (Roche 12013819001), 1:1,000 rabbit anti-caspase 8 (Cell Signaling Technology 4927), 1:1,000 rabbit anti-cleaved caspase 8 (Cell Signaling Technology 8592), 1:1,000 rabbit anti-MLKL(phospho Ser345, Abcam ab196436), 1:10,000 mouse anti-β actin (Sigma A2228) and 1:1,000 rabbit anti-HMGB1 (Abcam ab18256). Secondary antibodies used were 1:10,000 goat anti-rabbit HRP (Insight Biotechnology 474-1506) and 1:10,000 goat anti-mouse HRP (Insight Biotechnologies, 474–1806). HRP was detected using the Immobilon Western Chemiluminescent HRP substrate (Millipore), visualised on a ChemiDoc imaging system (BioRad) or a GE Amersham Imager 680.

### Restriction assays

#### High-content imaging

Following previously optimised protocols ^43,44^, MEFs or 75,000 BMDMs were seeded in black clear-bottom 96-well μ-Plate imaging plates (Ibidi 89626). Cells were pre-stimulated for 24 h with IFNγ according to host species: 100 U/ml mIFNγ (Thermo Fisher, Gibco #PMC4031) for murine BMDM and MEFs, and 25 U/ml mIFNγ for rat BMDM. Cells were infected at a MOI of 0.3 for 24 h, then fixed in 4% PFA and stained with 5 μg/ml DAPI and 5 μg/ml CellMask Deep Red (Invitrogen C10046). Plates were imaged using the Opera Phenix high-content screening system, with 25 images and 5 focal planes acquired per well, and an automated analysis of infection phenotypes was performed using Harmony v5 (PerkinElmer) as previously described ^44^. Data is reported as the mean proportion of each factor (vacuole number and total *Toxoplasma* number) in IFNγ-treated wells relative to untreated wells. Differences between strains were tested by paired two-sided *t* test with the Benjamini, Krieger and Yekutieli FDR correction.

#### Cytation5 plate reader

Cells were prepared, infected and analysed following the protocol above. As previously performed ^44^, plates were imaged on a Cytation5 plate reader (BioTek) using a 20x objective in a 2×2 tile. Since all strains express an mCherry reporter, the total signal was measured with the Texas Red filter (Ex/Em 586/647) and used as proxy of *Toxoplasma* growth.

#### Propidium Uptake assay

Black clear-bottom 96-well plates were seeded with 75,000 BMDMs per well and stimulated with 100 U/ml mIFNγ for 24 h. Cells were infected at an MOI of 3 in a media solution with 5 μg/mL propidium iodide (Thermo Fisher P3566). Images of each well were acquired every 30 minutes until 14 h post-infection in two different systems due to the lab relocation in a different institute. Similarly to previously established protocols ^43^, experiments in Figure 4A-B and Figure S4A-B, were analysed on a Nikon Ti-E inverted widefield fluorescence microscope with a Nikon CFI Plan Fluor 4x/0.13 objective and Hamamatsu C11440 ORCA Flash 4.0 camera running NIS Elements (Nikon). At the end of the experiment, 10% Triton X-100 solution was added to a final concentration of 1% v/v to fully permeabilise the cells and a final image was captured of each well. In each image, the total fluorescence signal was measured. The percentage of propidium iodide uptake in each well at each timepoint was calculated by subtracting the first measurement at 1 hpi to remove background fluorescence signal and normalising the total fluorescence intensity following full permeabilisation to 100% uptake. Experiments in Figure 4F-G and Figure S4F were analysed in an Agilent Cytation C5 plate reader with a 4x/0.13 objective and the Texas Red filter (Ex/Em 586/647). 30 minutes after infection at a MOI of 5, cells were washed with media with propidium iodide, with or without IFNγ based on the initial treatment, and with 1 μM ML10 (gift of Michael Blackman), or left untreated as control. All acquired images were analysed with the built-in software, and host cell nuclei positive to propidium iodide were automatically counted. Following Triton X-100 permeabilisation, another round of imaging was performed, and the number of dead cells was expressed as percentage of propidium iodide positive cells at each time point and condition out of the total in the final read. In both settings, imaging was performed at 37°C and with 5% CO2. All conditions were performed in technical triplicate, of which the mean was taken to represent each biological replicate. Differences between strains were tested at 9 hpi by One-Way ANOVA with paired two-sided *t*-test with the Benjamini, Krieger and Yekutieli FDR correction.

#### Transmission Electron Microscopy

1E5 PWD/Phj BMDM were seeded on 10 mm Poly-D-Lysine (Gibco A3890401) treated glass coverslips, treated for 24 h with 100 U/ml mIFNγ or left untreated, and infected with 5E5 parasites of the RH ΔKU80, ΔGRA12, ΔGRA12::GRA12-HA strains for 2 h. Cells were then washed 3x with PBS and fixed in 2.5% glutaraldehyde and 4% formaldehyde in 0.1 M phosphate buffer (PB; pH 7.4) for 30 min at room temperature. Samples were washed 2x with 0.1 M PB and stained with 1% (v/v) osmium tetroxide (Taab)/ 1.5% potassium ferricyanide (Sigma) for 1 hr at 4°C. All remaining processing steps were performed as per previously established protocols ^44^ using the Pelco BioWave Pro+ microwave (Ted Pella) with a Steady Temp set to 21°C. In brief, samples were incubated in 1% (w/v) tannic acid in 0.05 M PB (pH 7.4; Sigma) for 14 min under vacuum in 2 min cycles alternating with/without 100W power, followed by 1% sodium sulphate in 0.05 M PB (pH 7.4; Sigma) for 1 min without vacuum at 100 W. Samples were then washed in dH20 and dehydrated through an ethanol series (70%, 90% and 100%), followed by 3x incubations in 100% acetone at 250 W for 40 s without vacuum. Samples were exchanged into 1:1 acetone: Epon resin (Taab) overnight, then into 100% resin for 6 h before polymerising at 60°C for 48 h. The glass coverslip was removed with liquid nitrogen. Samples were sectioned using a UC7 ultramicrotome (Leica Microsystems) and 70 nm sections were picked up on Formvar-coated 2 mm copper slot grids (Gilder Grids). All grids were post stained with 3% lead citrate for 2 min. For all six conditions, sections containing *Toxoplasma gondii* in a horizontal orientation were viewed at 5000x, 6000x, 8000x, 10000x and 15000x using a 120 kV 1400FLASH TEM (JEOL), and images were acquired with a JEOL Matataki Flash sCMOS camera. The vacuolar area was quantified in Fiji ^72^ measuring the difference in area between the parasite membrane and the parasitophorous vacuole membrane (PVM). When the PVM was not visible, the vacuolar area was imputed to 0.01 μm^2^.

#### Griess Assay for Nitric Oxide quantification

75,000 C57BL/6J BMDMs were seeded in a 96 well plate and treated for 24 h with 100 U/ml IFNγ and 0.2 µg/ml LPS (InvivoGen 0111:B4), or left untreated as control before infection with RH ΔUPRT, ΔGRA12, ΔGRA12::GRA12-HA in a ratio cell:parasite of 1:5 and a total volume of 100 µl phenol-free media to not interfere with the colorimetric reaction. Uninfected cells were used as control and all conditions were performed in technical triplicate. After 24 h the plate was spun 1000 rpm for 1 min and 50 μl of supernatant were mixed in a 96 well plate in a 1:1 ratio with Griess reagent freshly prepared (1:1 solution of 0.1% N-(1-Naphthyl) ethylenediamine dihydrochloride (Sigma-Aldrich) in distilled water and 1% sulfanilamide (Sigma-Aldrich) in 5% phosphoric acid (Sigma-Aldrich)). A solution of sodium nitrite 0-100 μM was used to calibrate the standard curve. The plate was briefly shaken and incubated at room temperature for 5 min, then read at 540 nm in a Biotek Sinergy H1 Neo2 machine. To confirm the activity of the iNOS inhibitor N-(3-(Aminomethyl)benzyl)acetamidine (1400W-HCl, Selleck Chemicals S8337), 100 μM of the same were added 1 h before 0.2 µg/ml LPS stimulation of C57BL/6J BMDMs pretreated with 100 U/ml IFNγ or left untreated as control.

#### Immunoprecipitation and Mass Spectrometry

20E6 PWD/Phj BMDMs were seeded in 15 cm Petri dishes and treated for 24 h with 10 U/ml mIFNγ before infection with RH ΔKU80 or RH GRA12-HA in triplicate for 24 h at a MOI 0.3. Infected cells were washed 2x in cold PBS then lysed in cold immunoprecipitation (IP) buffer (10 mM Tris, 150 mM NaCL, 0.5 mM EDTA and 0.4% NP40, pH 7.5 in H2O, supplemented with 2x cOmplete Mini EDTA-free Protease Inhibitor Cocktail). Lysates were syringe-lysed 6x through a 30G needle and left on ice for 1 h, then centrifuged at 2,000 *g* for 20 min to remove the insoluble fraction. Soluble fractions were added to 30 μl/sample anti-HA agarose beads (Thermo 26182), then incubated o/n at 4°C with rotation. Beads were washed 3x with cold IP buffer for 10 min each, then proteins were eluted in 30 μl 4x Sample Loading Buffer supplemented with DTT and boiled for 5 min at 95°C.

Approximately 20 μl of each IP elution was loaded on a 10% Bis-Tris gel and run into the gel for 1 cm, then stained with InstantBlue Coomassie Protein Stain (Abcam ab119211). Proteins were alkylated in-gel prior to digestion with 100 ng trypsin (modified sequencing grade, Promega) overnight at 37°C. Supernatants were dried in a vacuum centrifuge and resuspended in 0.1% trifluoroacetic acid (TFA), and 1 to 10 μl of acidified protein digest was loaded onto a 20 mm × 75 μm Pepmap C18 trap column (Thermo Scientific) on an Ultimate 3000 nanoRSLC HPLC (Thermo Scientific) prior to elution via a 50 cm × 75 μm EasySpray C18 column into a Lumos Tribrid Orbitrap mass spectrometer (Thermo Scientific). A 70’ gradient of 6% to 40% B was used to elute bound peptides followed by washing and re-equilibration (A = 0.1% formic acid, 5% DMSO; B = 80% ACN, 5% DMSO, 0.1% formic acid). The Orbitrap was operated in “Data Dependent Acquisition” mode followed by MS/MS in “TopS” mode using the vendor supplied “universal method” with default parameters.

Raw files were processed to identify tryptic peptides using Maxquant (maxquant.org) and searched against the *Toxoplasma* (ToxoDB-56_TgondiiGT1_AnnotatedProteins) and Murine (Uniprot, UP000000589) reference proteome databases and a common contaminants database. A decoy database of reversed sequences was used to filter false positives, at peptide and protein false detection rates (FDRs) of 1%. *T* test-based volcano plots of fold changes were generated in Perseus (<u>maxquant.net/perseus</u>) with significantly different changes in protein abundance determined by a permutation-based FDR of 0.05% to address multiple hypothesis testing. Raw data are provided in Table S8.

#### Phylogenetic analysis of GRA12-like proteins and protein structure prediction

Amino acid sequences of homologues and paralogues of GRA12 within Apicomplexa were obtained from the ToxoDB database ^73^. A global alignment of the 25 sequences was performed in Geneious Prime (version 2024.0.2. Cost matrix Blosum62, gap open penalty 12, gap extension penalty 3 and refinement iterations 2). A Neighbor-Joining Tree was built with Bootstrap method with 100 replicates and imputing GRA12D (TGGT1_308970) as outgroup.

GRA12’s structures from *Toxoplasma* (UniProt S7VY87) and *Neospora* (A0A0F7UEK3) as predicted by the AlphaFold protein structure database (version 3) ^74^, were overlapped on PyMOL (version 2.4.1), and the Root Mean Square Deviation (RMSD) was calculated to infer protein structure similarity. The C and N termini were not included in Figure 5C.

## Acknowledgements

We thank the Science Technology Platforms at the Francis Crick Institute – and in particular the Biology Research Facility, the Advanced Sequencing Facility, the Light Microscopy Facility and the Cell Services Facility – for their support and help. We thank the Advanced Imaging Facility and Genomics Facility at the Instituto Gulbenkian de Ciência for their support and help. We thank Eva Frickel for the VAND strain and Martin Blume for the VEG strain. We thank James Turner for the PWD/Phj and CAST/Eij strains, and Sangrithi Mahesh for CAST/Eij tissue samples. We thank Andreas Wack for providing the BMDMs differentiation protocol, Jonathan Howard, Maryse Lebrun, David Sibley and Jean François Dubremetz for providing antibodies, Michael Blackman for providing reagents and the Soares lab for providing the L929 cell line. We thank VEuPathDB for providing access to the *Toxoplasma* databases ^73^. We thank all members of the Treeck lab, Max Gutierrez, Beren Aylan and Alice Balard for critical input and reading of this manuscript. This work was supported by an award to M.T. from the Wellcome Trust (223192/Z/21/Z) and by funding to M.T. from the Francis Crick Institute which receives its core funding from Cancer Research UK, the UK Medical Research Council and the Wellcome Trust (CC2132, CC0199 and CR2023/030/2123). F.T. is supported by the Deutsche Forschungsgemeinschaft (TO 1349/1-1).

## Author contributions

Conceptualization, F.T. and M.T.; methodology, F.T., S.B. and E.L.; investigation, F.T. S.B., E.L., O.S. and J.P.F.; formal analysis, F.T and S.B.; visualisation, F.T.; writing – original draft, F.T and M.T.; writing – review and editing, all authors; supervision, M.T.; funding acquisition, F.T. and M.T.

## Supplementary Information

### Supplementary Figures

Combined file with supplementary figures and figure legends.

**Figure S1.**
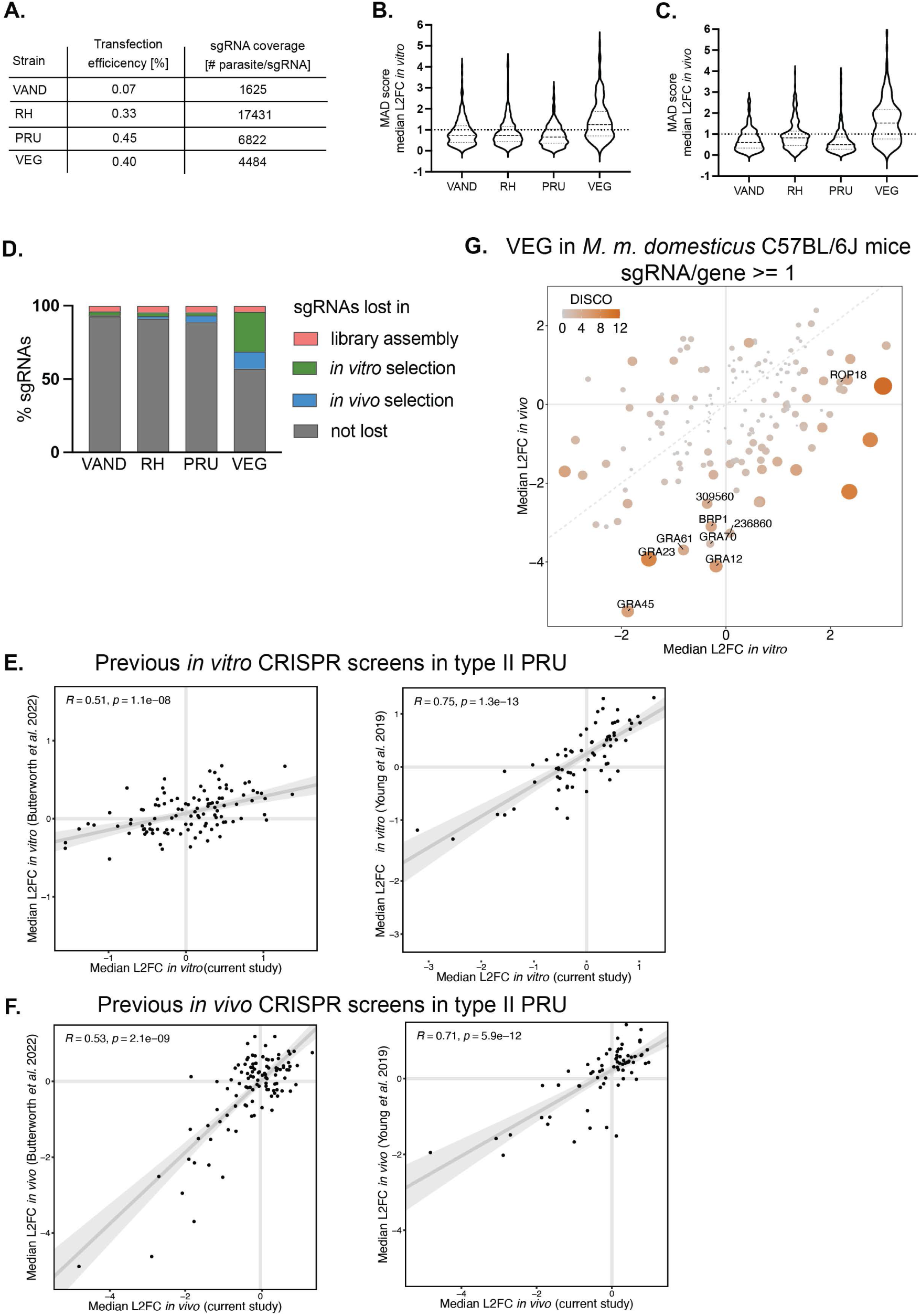
*In vivo* CRISPR screens of the secretome of four *Toxoplasma* strains. (A) Transfection efficiency and relative sgRNA coverage in each screen. (B) Median Absolute Deviation (MAD) of the log2 fold change (L2FC) of sgRNA read counts between the plasmid and the inoculum and (C) of the inoculum and the mouse peritoneum in each screen. (D) sgRNA loss at each screen step. (E) Correlation of the median L2FC *in vitro* and (F) *in vivo* of the current type II screen with previous screens performed in our research group ^42,43^. (G) Scatter plots of the median L2FC for each gene *in vitro* and *in vivo* of the CRISPR screen of VEG in C57BL/6J mice. The colour and size of each point reflects the Discordance/Concordance (DISCO) score, and the dashed grey line indicates equal L2FC. A threshold of 1 sgRNA per gene was applied compared to 3 sgRNAs per gene in Figure 1B.

**Figure S2.**
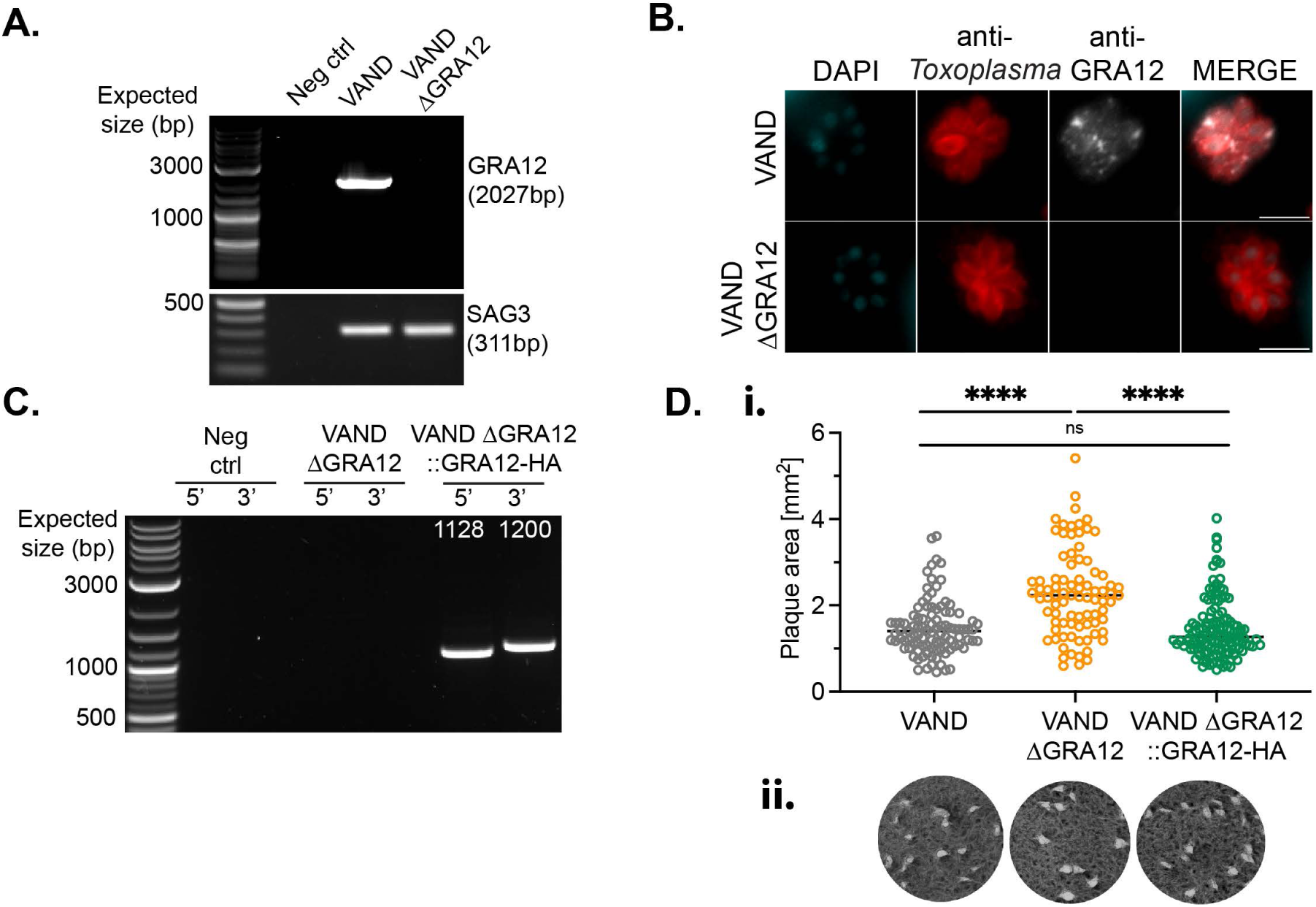
Establishment of VAND ΔGRA12 and ΔGRA12::GRA12-HA strains. (A) PCR amplification of the GRA12 locus in the parental VAND and ΔGRA12 strains. Amplification of SAG3 was performed as control. (B) Immunofluorescence verification of GRA12 in the VAND ΔGRA12 strain. Scale bar represents 10 µm. (C) PCR validation of the VAND ΔGRA12::GRA12-HA strain. (D) Scatter plot of the plaque size of VAND parental and derived clones, bar represents the median. Significance was tested using the one-way ANOVA test, N=2 (i) and respective images (ii).

**Figure S3.**
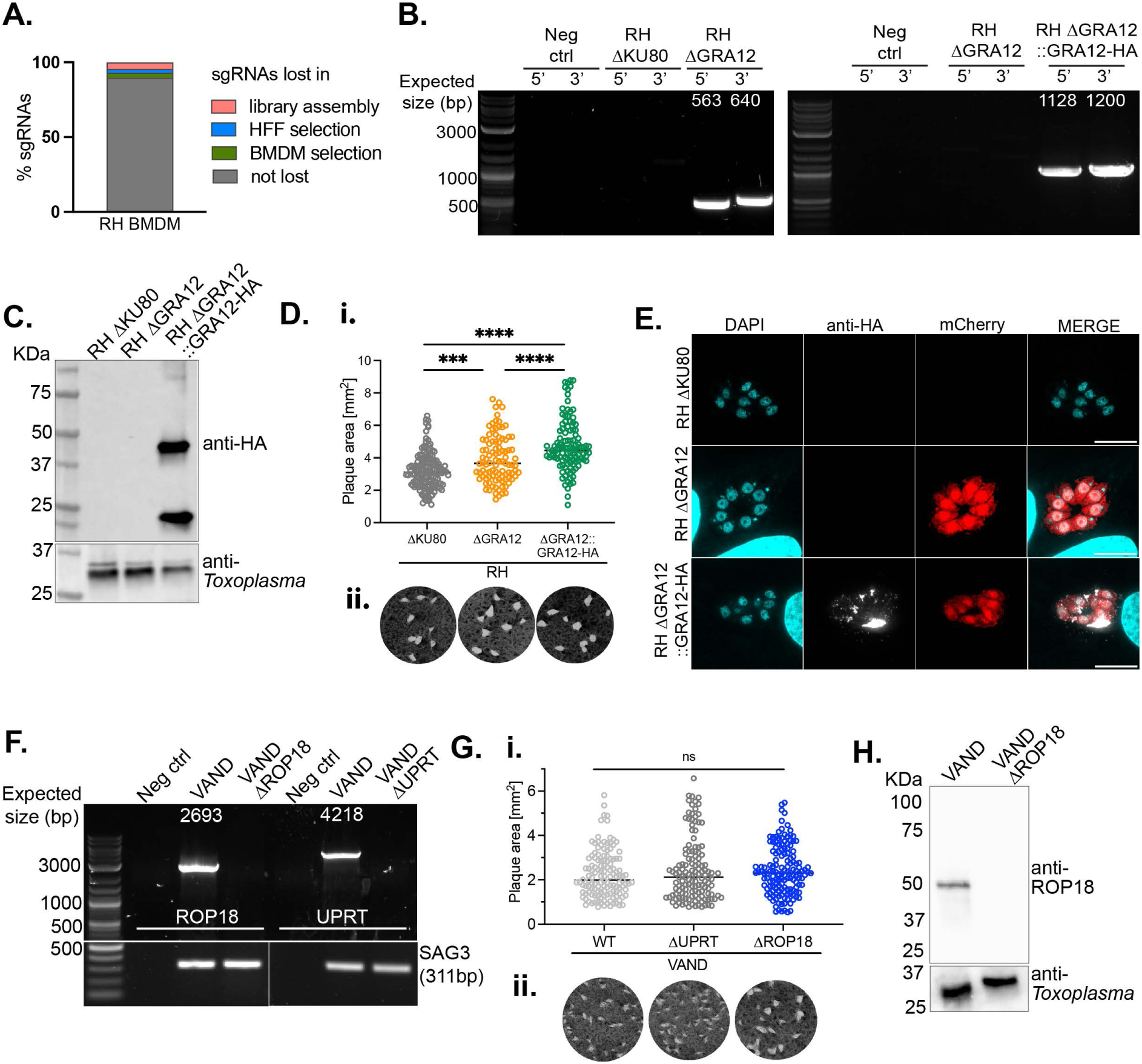
Creation of RH and VAND KO mutants for the validation of GRA12 as transcendent virulence factor *in vitro*. (A) sgRNA loss at each screen step of the type I RH CRISPR screen in BMDMs. (B) PCR validation of the RH ΔGRA12 and complemented strains. (C) Verification of the GRA12::HA expression via anti-HA detection in the RH ΔGRA12::GRA12-HA strain by western blot. (D) Plaque size of RH ΔKU80 parental and derived clones. Significance was tested using a one-way ANOVA, N=2, p *** <0.001, *** <0.0001 (i), and respective images (ii). (E) Immunofluorescence verification of the C-terminal HA-tagged GRA12 in the RH ΔGRA12::GRA12-HA strain. Scale bar is 10 µm. (F) PCR validation of the VAND ΔUPRT and ΔROP18. Amplification of SAG3 was performed as control. (G) Plaque size of VAND parental and derived clones ΔUPRT and ΔROP18. Significance was tested using an unpaired t test, N=1 (i), and respective images (ii). (H) Verification of the ROP18 KO via anti-ROP18 detection in the VAND ΔROP18 strain by western blot.

**Figure S4.**
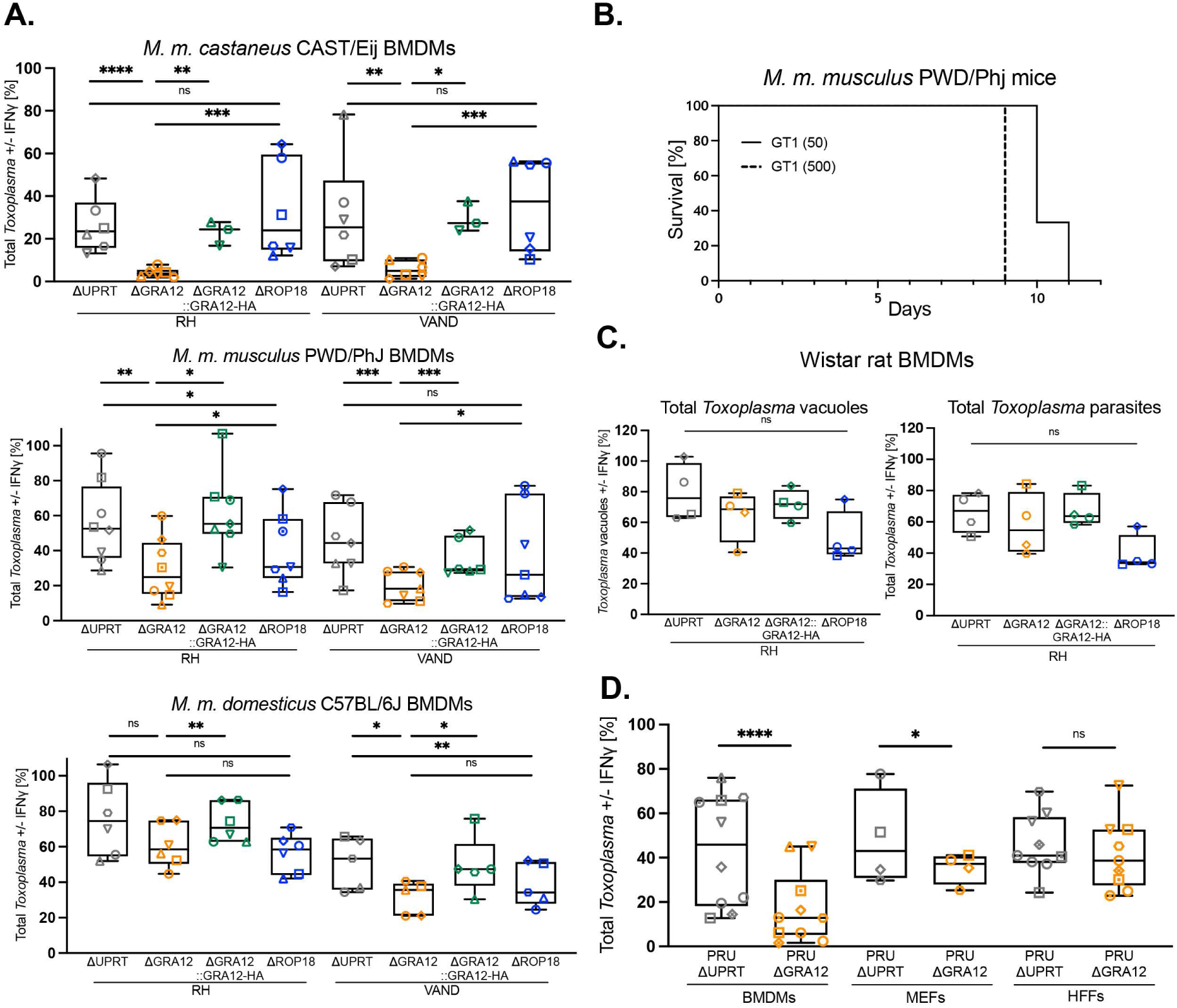
Validation of GRA12 as *Toxoplasma* transcendent virulence factor in mouse subspecies. (A) Quantification of high content-automated imaging of parasite numbers in IFNγ-treated BMDMs of different mouse subspecies relative to untreated controls. Cells were infected with RH ΔUPRT, ΔGRA12, ΔGRA12::GRA12-HA and ΔROP18 and *Toxoplasma* parasites were quantified at 24 h after infection. (B) Survival curve of PWD/Phj mice infected with the type I GT1 strain, dose in parenthesis. N=3 mice per group. (C) Quantification of high-content automated imaging of *Toxoplasma* infection in IFNγ-treated Wistar rat BMDMs or (D) murine embryonic fibroblasts (MEFs), human foreskin fibroblasts (HFFs) and murine BMDMs, relative to untreated controls. Cells were infected with RH ΔUPRT, ΔGRA12, ΔGRA12::GRA12-HA and ΔROP18, or PRU ΔUPRT and PRU ΔGRA12, and parasites were quantified at 24 h after infection. Symbol shapes indicate biological repeats. Significance was tested using the One-way Anova test with the Benjamini, Krieger and Yekutieli FDR correction. p * <0.05, ** <0.01, *** <0.001, **** <0.0001.

**Figure S5.**
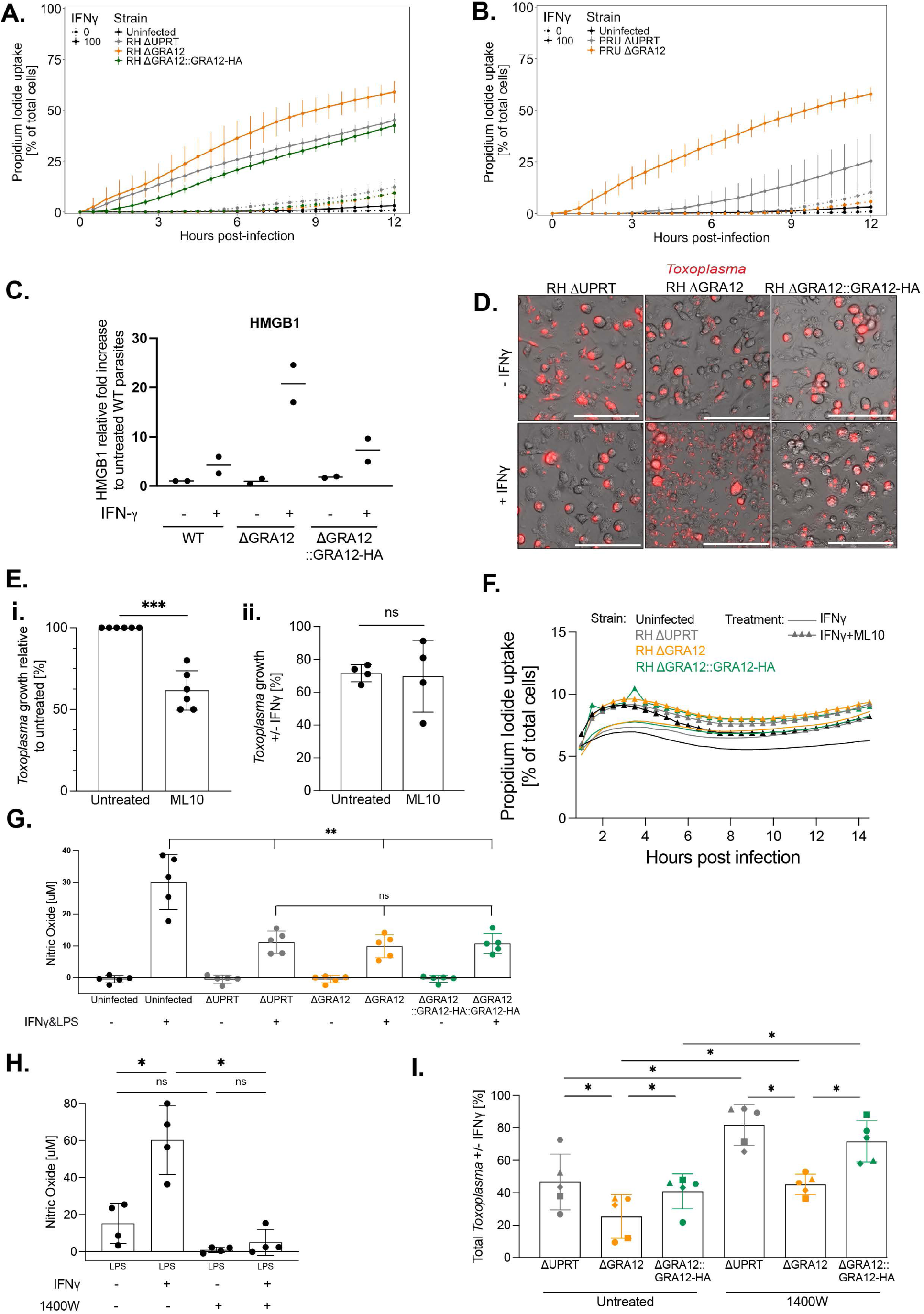
Lack of GRA12 results in IFNγ-mediated host cell death by necrosis. Time course of PWD/Phj BMDMs cell death quantified via Propidium Iodide uptake. Cells were pre-stimulated for 24 h with IFNγ (continue line) or left untreated (dashed line) and then infected with (A) RH ΔUPRT, ΔGRA12, ΔGRA12::GRA12-HA or (B) PRU ΔUPRT or ΔGRA12, or left uninfected as a control. The number of dead cells is expressed as a percentage of the total at each time point. (C) Quantification of the HMGB1 necrosis marker from the western blot analysis of the supernatant of infected BMDMs, relative to the untreated parental condition. (D) Microscopy images of IFNγ-treated or untreated C57BL/6J BMDMs, infected for 24 h with RH ΔUPRT, ΔGRA12 or ΔGRA12::GRA12-HA. Scale bar represents 100 µm. (E) Quantification by imaging of *Toxoplasma* growth in ML10-treated relative to untreated BMDMs, in naïve conditions (i) or in IFNγ-treated conditions relative to untreated controls (ii). (F) Time course of C57BL/6J BMDMs cell death quantified via Propidium Iodide uptake. Cells were infected with RH ΔUPRT, ΔGRA12, ΔGRA12::GRA12-HA or left uninfected, and 1 hpi treated with 1 µM ML10 (triangle symbol) or left untreated as a control (continue line). The number of dead cells is expressed as a percentage of the total at each time point. (G) Quantification of Nitric Oxide from C57BL/6J BMDMs stimulated for 24 h with 100 U/ml IFNγ and 0.2 µg/ml LPS or left untreated as control, and infected for 24 h with RH ΔUPRT, ΔGRA12, ΔGRA12::GRA12-HA or left uninfected. Significance was tested using the One-way Anova test with the Benjamini, Krieger and Yekutieli FDR correction, (H) Quantification of Nitric Oxide from a culture of uninfected C57BL/6J BMDMs pretreated for 24 h with 100 U/ml IFNγ and with the iNOS inhibitor 1400W, or left untreated as control, and stimulated for 24 h with 0.2 µg/ml LPS. Significance was tested using the One-way Anova test with the Benjamini, Krieger and Yekutieli FDR correction, (I) Quantification by imaging of *Toxoplasma* restriction in IFNγ-treated or untreated C57BL/6 BMDMs, pretreated of not with 1400W, infected with RH ΔUPRT, ΔGRA12, ΔGRA12::GRA12-HA and parasites were quantified at 24 h after infection. Symbol shapes indicate biological repeats. P *<0.05, ** < 0.01.

**Figure S6.**
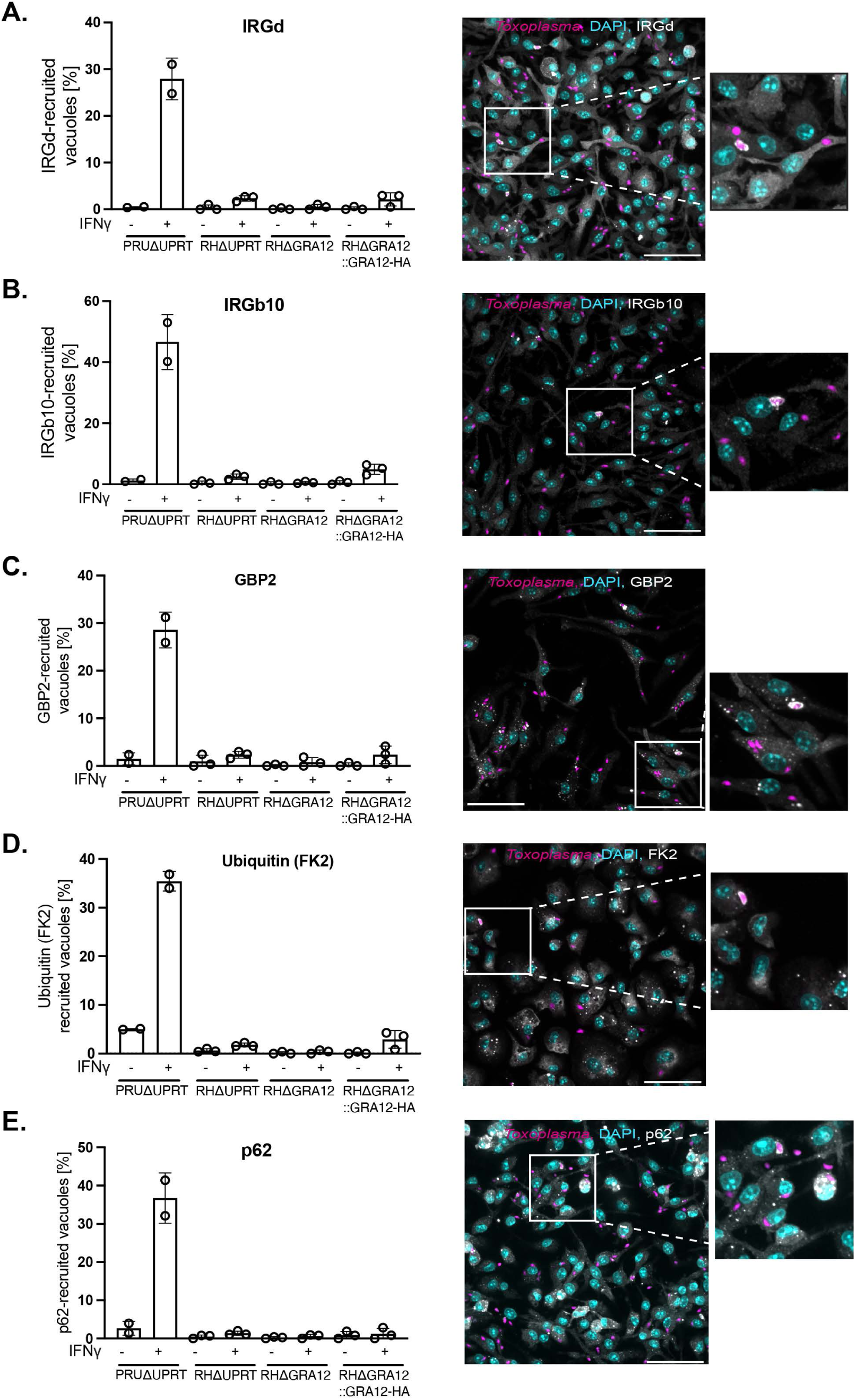
Recruitment of host factors to the PVM. Quantification of the recruitment to the PVM of host proteins: IRGd (A), IRGb10 (B), GBP2 (C), Ubiquitin (FK2, D) and p62 (E) with representative images on the right panels. C57BL/6J BMDMs were pretreated for 24h with IFNγ or left untreated, and infected for 90 minutes with RH ΔUPRT, ΔGRA12 or ΔGRA12::GRA12-HA, before being fixed and stained for immunofluorescence. N=3 and N=2 for the PRU ΔUPRT line used as positive control. Scale bar represents 50 µm.

**Figure S7.**
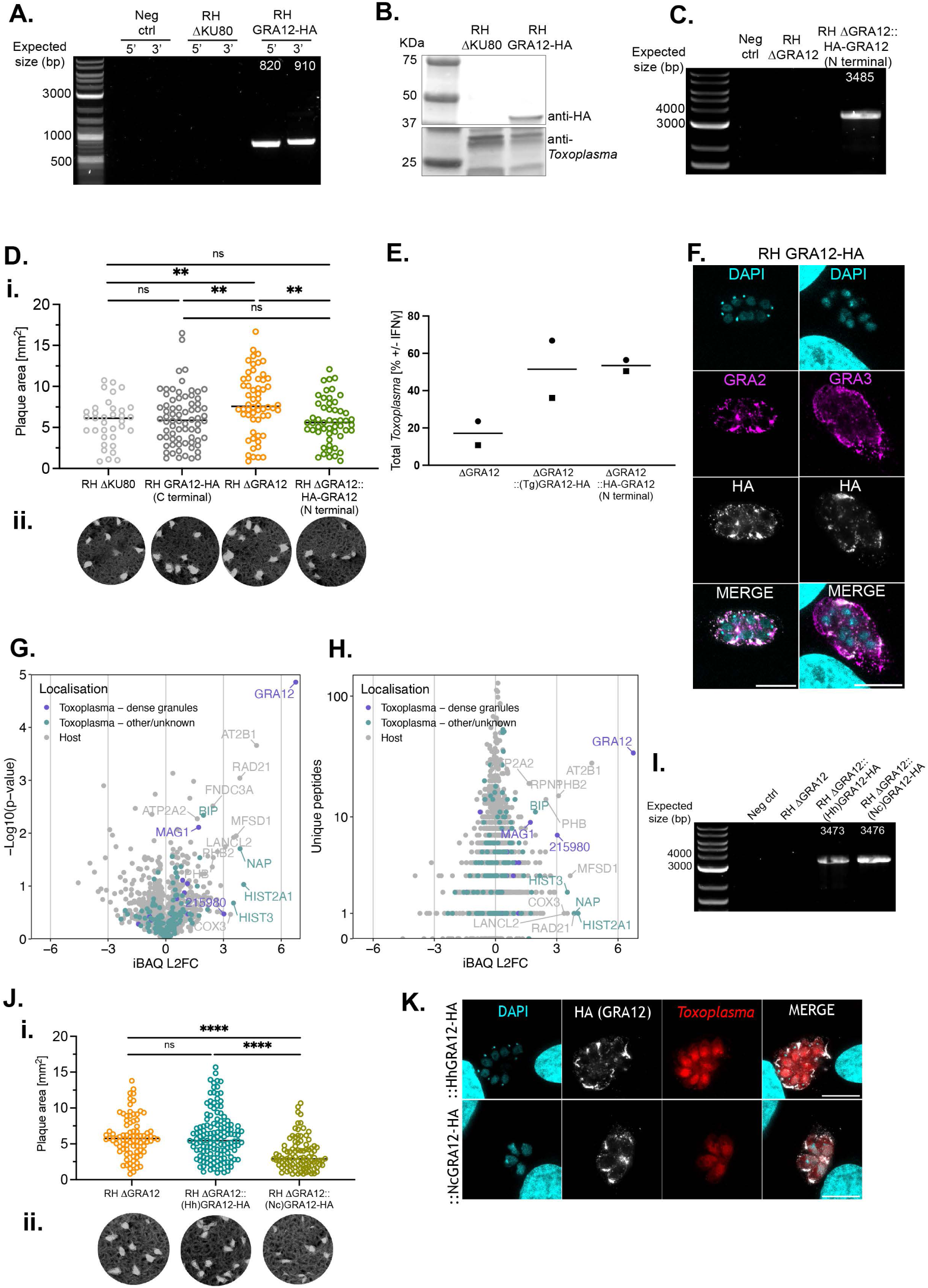
Creation of the N and C terminally HA-tagged GRA12 strains and validation of GRA12 paralogs activity. (A) PCR validation of the RH GRA12 C-terminally HA-tagged line in the endogenous locus (RH GRA12-HA) by PCR and (B) by western blot. (C) PCR validation of the RH GRA12 N-terminally HA-tagged line (RH ΔGRA12::HA-GRA12) created by complementing GRA12 in the *Uprt* locus. (D) Plaque size of RH GRA12-HA and RH ΔGRA12::HA-GRA12 lines compared to their respective parental RH ΔKU80 and RH ΔGRA12 strains. Significance was tested using an unpaired t test, N=1 (i) and respective images (ii). (E) Relative *Toxoplasma* growth in IFNγ-treated versus untreated BMDMs. BMDMs were infected with the RH ΔGRA12 strain, or the RHΔGRA12 strain complemented with *Toxoplasma* (Tg) GRA12 or the N-terminally HA-tagged line RH ΔGRA12::HA-GRA12, or the parental RH ΔGRA12 strain as control for 24h before a plate reader quantification of the mCherry signal as proxy for parasite growth. N=2. (F) Immunofluorescence localisation of GRA2 (left panel) and GRA3 (right panel) in relation to GRA12 in the RH GRA12-HA line infecting HFFs. Scale bar represents 10 µm. (G) Volcano plot of proteins identified by mass spectrometry in the anti-HA pull down in lysates from IFNγ-treated BMDM infected with either RH GRA12-HA or the parental RH ΔKU80 strains. The relative abundance of the proteins (iBAQ L2FC) is plotted against the p-value (-Log10(p-value), (G)) and against the number of peptides detected ((Unique peptides), (H)). (I) PCR validation of the RH ΔGRA12 strain complemented with the *H. hammondi* (::(Hh)GRA12-HA) or the *N. caninum* (::(Nc)GRA12-HA) GRA12 homologues. (J) Plaque size of the complemented::(Hh)GRA12-HA and::(Nc)GRA12-HA strains, compared to the parental RH ΔGRA12 strain. Significance was tested using an unpaired t test, N=1 (i) and respective images (ii). (K) Immunofluorescence localisation of GRA12 via anti-HA in the RH ΔGRA12::(Hh)GRA12-HA strain (upper panel) and RH ΔGRA12::(Nc)GRA12-HA strain (lower panel). Scale bar represents 10 µm. p ** <0.01, **** <0.0001.

**Figure S8.**
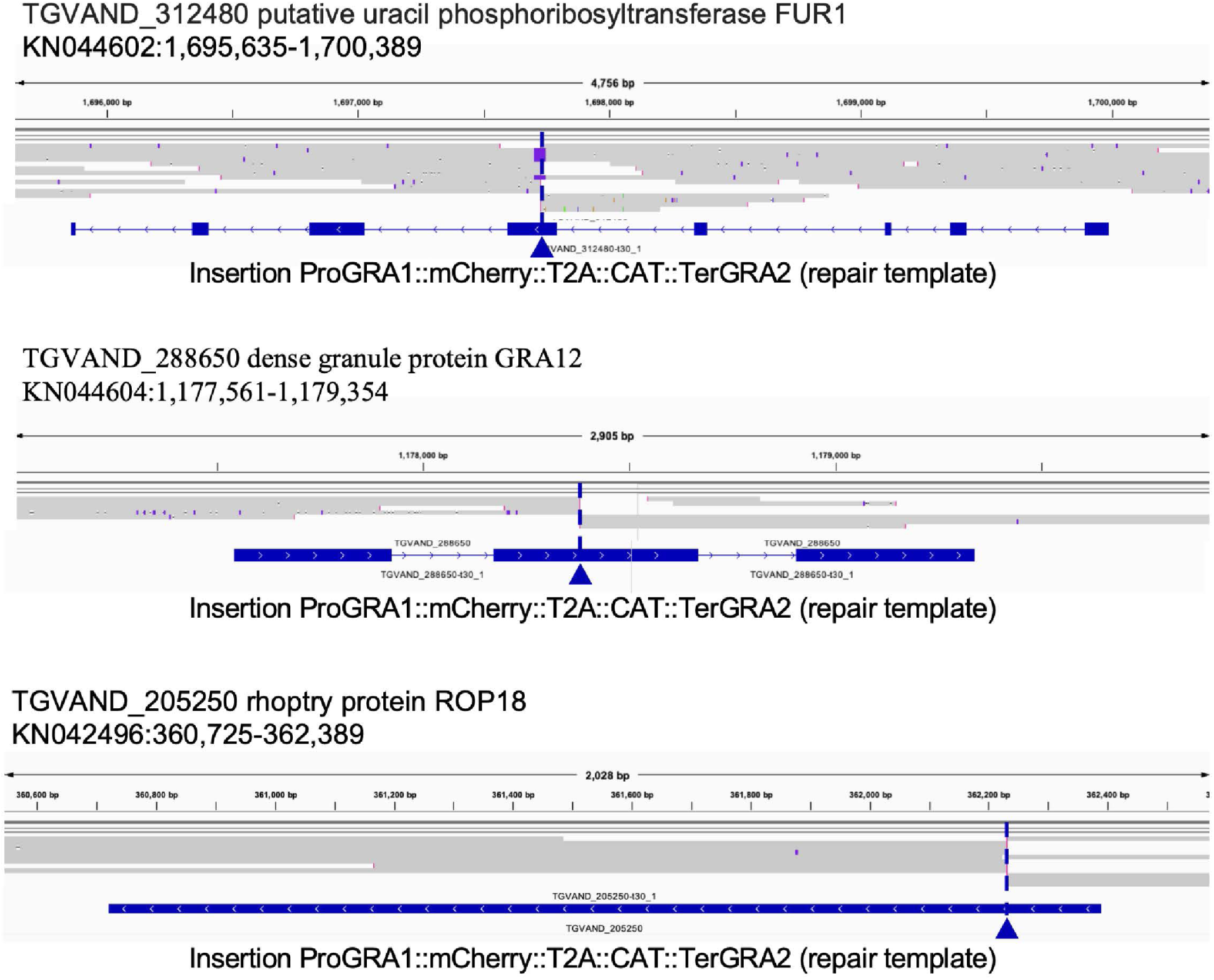
Validation by Nanopore sequencing of the VAND mutant strains. Alignment of reads from Nanopore sequencing to the reference genome assembly ToxoDB-65_TgondiVAND. The alignment shows disruption of the endogenous UPRT (TGVAND_312480, upper panel). GRA12 (TGVAND_205250, middle panel) and ROP18 (TGVAND_205250, lower panel) loci, for the creation of the VAND ΔUPRT, ΔGRA12 and ΔROP18 strains respectively. The arrowhead the integration of the repair template PriGRA1::McGarry::T2A::CAT::TerGRA2 in the coding sequence.

---

**Supplementary Movie S1:** Brightfield movie of IFNγ-treated BMDMs infected with RH ΔGRA12 parasites.

**Table S1.** List of primers used in the study.

**Table S2. VAND CRISPR knockout screen results.** A) Raw protospacer sequencing read counts. B) Normalised protospacer sequencing read counts. C) Protospacer L2FCs *in vivo* and *in vitro*. D) Gene L2FCs *in vivo*, *in vitro* and their difference, p-values, L2FCs median absolute deviation (MAD) and DISCO scores.

**Table S3. RH CRISPR knockout screen results.** A) Raw protospacer sequencing read counts. B) Normalised protospacer sequencing read counts. C) Protospacer L2FCs *in vivo* and *in vitro*. D) Gene L2FCs *in vivo*, *in vitro* and their difference, p-values, L2FCs median absolute deviation (MAD) and DISCO scores.

**Table S4. PRU CRISPR knockout screen results.** A) Raw protospacer sequencing read counts. B) Normalised protospacer sequencing read counts. C) Protospacer L2FCs *in vivo* and *in vitro*. D) Gene L2FCs *in vivo*, *in vitro* and their difference, p-values, L2FCs median absolute deviation (MAD) and DISCO scores.

**Table S5. VEG CRISPR knockout screen results.** A) Raw protospacer sequencing read counts. B) Normalised protospacer sequencing read counts. C) Protospacer L2FCs *in vivo* and *in vitro*. D) Gene L2FCs *in vivo*, *in vitro* and their difference, p-values, L2FCs median absolute deviation (MAD) and DISCO scores.

**Table S6. Gene L2FCs and their difference in all screens.** A) Gene L2FCs *in vivo*, *in vitro* and their difference, p-values, L2FCs median absolute deviation (MAD) and DISCO scores. Genes are ranked for each screen based on the L2FC difference in vivo vs in vitro. B) Top ranking genes shared between the VAND, RH and PRU screens.

**Table S7. RH CRISPR knockout screen results in BMDMs.** A) Raw protospacer sequencing read counts. B) Normalised protospacer sequencing read counts. C) Protospacer L2FCs in IFNγ-treated versus untreated BMDMs. D) Gene L2FCs in IFNγ-treated BMDMs, in untreated BMDMs and their difference, p-values, L2FCs median absolute deviation (MAD) and DISCO scores.

**Table S8. GRA12 co-immunoprecipitation mass spectrometry data.**

## Bibliography

1. Pappas, G., Roussos, N. & Falagas, M. E. Toxoplasmosis snapshots: global status of Toxoplasma gondii seroprevalence and implications for pregnancy and congenital toxoplasmosis. Int. J. Parasitol. 39, 1385–1394 (2009).

2. Su, C., Howe, D. K., Dubey, J. P., Ajioka, J. W. & Sibley, L. D. Identification of quantitative trait loci controlling acute virulence in Toxoplasma gondii. Proc. Natl. Acad. Sci. U. S. A. 99, 10753–10758 (2002).

3. Galal, L., Hamidović, A., Dardé, M. L. & Mercier, M. Diversity of Toxoplasma gondii strains at the global level and its determinants. Food Waterborne Parasitol. 15, e00052 (2019).

4. Howe, D. K. & Sibley, L. D. Toxoplasma gondii comprises three clonal lineages: correlation of parasite genotype with human disease. J. Infect. Dis. 172, 1561–1566 (1995).

5. Grigg, M. E., Ganatra, J., Boothroyd, J. C. & Margolis, T. P. Unusual abundance of atypical strains associated with human ocular toxoplasmosis. J. Infect. Dis. 184, 633–639 (2001).

6. Shwab, E. K. et al. Geographical patterns of Toxoplasma gondii genetic diversity revealed by multilocus PCR-RFLP genotyping. Parasitology 141, 453–461 (2014).

7. Dardé, M. L. Toxoplasma gondii, “new” genotypes and virulence. Parasite 15, 366–371 (2008).

8. Dubey, J. P. Outbreaks of clinical toxoplasmosis in humans: five decades of personal experience, perspectives and lessons learned. Parasit. Vectors 14, 263 (2021).

9. Jensen, K. D. C. et al. Toxoplasma gondii superinfection and virulence during secondary infection correlate with the exact ROP5/ROP18 allelic combination. mBio 6, e02280 (2015).

10. Hakimi, M.-A., Olias, P. & Sibley, L. D. Toxoplasma Effectors Targeting Host Signaling and Transcription. Clin. Microbiol. Rev. 30, 615–645 (2017).

11. Olias, P., Etheridge, R. D., Zhang, Y., Holtzman, M. J. & Sibley, L. D. Toxoplasma Effector Recruits the Mi-2/NuRD Complex to Repress STAT1 Transcription and Block IFN-γ-Dependent Gene Expression. Cell Host Microbe 20, 72–82 (2016).

12. Gay, G. et al. Toxoplasma gondii TgIST co-opts host chromatin repressors dampening STAT1-dependent gene regulation and IFN-γ-mediated host defenses. J. Exp. Med. 213, 1779–1798 (2016).

13. Matta, S. K. et al. Toxoplasma gondii effector TgIST blocks type I interferon signaling to promote infection. Proc. Natl. Acad. Sci. 116, 17480–17491 (2019).

14. Saeij, J. P. J. et al. Toxoplasma co-opts host gene expression by injection of a polymorphic kinase homologue. Nature 445, 324–327 (2007).

15. Jensen, K. D. C. et al. Toxoplasma Polymorphic Effectors Determine Macrophage Polarization and Intestinal Inflammation. Cell Host Microbe 9, 472–483 (2011).

16. Barylyuk, K. et al. A Comprehensive Subcellular Atlas of the Toxoplasma Proteome via hyperLOPIT Provides Spatial Context for Protein Functions. Cell Host Microbe 28, 752–766.e9 (2020).

17. Reese, M. L., Zeiner, G. M., Saeij, J. P. J., Boothroyd, J. C. & Boyle, J. P. Polymorphic family of injected pseudokinases is paramount in Toxoplasma virulence. Proc. Natl. Acad. Sci. U. S. A. 108, 9625–9630 (2011).

18. Taylor, S. et al. A Secreted Serine-Threonine Kinase Determines Virulence in the Eukaryotic Pathogen Toxoplasma gondii. Science 314, 1776–1780 (2006).

19. Steinfeldt, T. et al. Phosphorylation of Mouse Immunity-Related GTPase (IRG) Resistance Proteins Is an Evasion Strategy for Virulent Toxoplasma gondii. PLOS Biol. 8, e1000576 (2010).

20. Bekpen, C. et al. The interferon-inducible p47 (IRG) GTPases in vertebrates: loss of the cell autonomous resistance mechanism in the human lineage. Genome Biol. 6, R92 (2005).

21. Gazzinelli, R. T., Mendonça-Neto, R., Lilue, J., Howard, J. & Sher, A. Innate resistance against Toxoplasma gondii: an evolutionary tale of mice, cats, and men. Cell Host Microbe 15, 132–138 (2014).

22. da Ferreira-da-Silva, M. F., da Fonseca Ferreira-da-Silva, M., Springer-Frauenhoff, H. M., Bohne, W. & Howard, J. C. Identification of the microsporidian Encephalitozoon cuniculi as a new target of the IFNγ-inducible IRG resistance system. PLoS Pathog. 10, e1004449 (2014).

23. Hunn, J. P., Feng, C. G., Sher, A. & Howard, J. C. The immunity-related GTPases in mammals: a fast-evolving cell-autonomous resistance system against intracellular pathogens. Mamm. Genome 22, 43–54 (2011).

24. Khaminets, A. et al. Coordinated loading of IRG resistance GTPases on to the Toxoplasma gondii parasitophorous vacuole. Cell. Microbiol. 12, 939–961 (2010).

25. Maric-Biresev, J. et al. Loss of the interferon-γ-inducible regulatory immunity-related GTPase (IRG), Irgm1, causes activation of effector IRG proteins on lysosomes, damaging lysosomal function and predicting the dramatic susceptibility of Irgm1-deficient mice to infection. BMC Biol. 14, 33 (2016).

26. Haldar, A. K. et al. IRG and GBP Host Resistance Factors Target Aberrant, “Non-self” Vacuoles Characterized by the Missing of “Self” IRGM Proteins. PLOS Pathog. 9, e1003414 (2013).

27. Lee, Y. et al. p62 Plays a Specific Role in Interferon-γ-Induced Presentation of a Toxoplasma Vacuolar Antigen. Cell Rep. 13, 223–233 (2015).

28. Dockterman, J. & Coers, J. How did we get here? Insights into mechanisms of immunity-related GTPase targeting to intracellular pathogens. Curr. Opin. Microbiol. 69, 102189 (2022).

29. Selleck, E. M. et al. Guanylate-binding Protein 1 (Gbp1) Contributes to Cell-autonomous Immunity against Toxoplasma gondii. PLOS Pathog. 9, e1003320 (2013).

30. Kravets, E. et al. Guanylate binding proteins directly attack Toxoplasma gondii via supramolecular complexes. eLife 5, e11479.

31. Fisch, D., Clough, B., Khan, R., Healy, L. & Frickel, E.-M. Toxoplasma-proximal and distal control by GBPs in human macrophages. Pathog. Dis. 79, ftab058 (2021).

32. Zhao, X.-Y. et al. iNOS is necessary for GBP-mediated T. gondii clearance in murine macrophages via vacuole nitration and intravacuolar network collapse. Nat. Commun. 15, 2698 (2024).

33. Lilue, J., Müller, U. B., Steinfeldt, T. & Howard, J. C. Reciprocal virulence and resistance polymorphism in the relationship between Toxoplasma gondii and the house mouse. eLife 2, e01298 (2013).

34. Niedelman, W., Sprokholt, J. K., Clough, B., Frickel, E.-M. & Saeij, J. P. J. Cell death of gamma interferon-stimulated human fibroblasts upon Toxoplasma gondii infection induces early parasite egress and limits parasite replication. Infect. Immun. 81, 4341– 4349 (2013).

35. Fisch, D. et al. Human GBP1 is a microbe-specific gatekeeper of macrophage apoptosis and pyroptosis. EMBO J. 38, e100926 (2019).

36. Zhao, Y. O., Khaminets, A., Hunn, J. P. & Howard, J. C. Disruption of the Toxoplasma gondii parasitophorous vacuole by IFNgamma-inducible immunity-related GTPases (IRG proteins) triggers necrotic cell death. PLoS Pathog. 5, e1000288 (2009).

37. Khan, A., Taylor, S., Ajioka, J. W., Rosenthal, B. M. & Sibley, L. D. Selection at a Single Locus Leads to Widespread Expansion of Toxoplasma gondii Lineages That Are Virulent in Mice. PLOS Genet. 5, e1000404 (2009).

38. Murillo-León, M. et al. Molecular mechanism for the control of virulent Toxoplasma gondii infections in wild-derived mice. Nat. Commun. 10, 1233 (2019).

39. Hassan, M. A., Olijnik, A.-A., Frickel, E.-M. & Saeij, J. P. Clonal and atypical Toxoplasma strain differences in virulence vary with mouse sub-species. Int. J. Parasitol. 49, 63–70 (2019).

40. English, E. D., Adomako-Ankomah, Y. & Boyle, J. P. Secreted effectors in Toxoplasma gondii and related species: determinants of host range and pathogenesis? Parasite Immunol. 37, 127–140 (2015).

41. Ehret, T., Torelli, F., Klotz, C., Pedersen, A. B. & Seeber, F. Translational Rodent Models for Research on Parasitic Protozoa-A Review of Confounders and Possibilities. Front. Cell. Infect. Microbiol. 7, 238 (2017).

42. Young, J. et al. A CRISPR platform for targeted in vivo screens identifies Toxoplasma gondii virulence factors in mice. Nat. Commun. 10, 3963 (2019).

43. Butterworth, S. et al. Toxoplasma gondii virulence factor ROP1 reduces parasite susceptibility to murine and human innate immune restriction. PLoS Pathog. 18, e1011021 (2022).

44. Lockyer, E. J. et al. A heterotrimeric complex of Toxoplasma proteins promotes parasite survival in interferon gamma stimulated human cells. 2022.12.08.519568 Preprint at 10.1101/2022.12.08.519568 (2022).

45. Christiansen, C. et al. In vitro maturation of Toxoplasma gondii bradyzoites in human myotubes and their metabolomic characterization. Nat. Commun. 13, 1168 (2022).

46. Sidik, S. M. et al. A Genome-Wide CRISPR Screen in Toxoplasma Identifies Essential Apicomplexan Genes. Cell 166, 1423–1435.e12 (2016).

47. Wang, Y. et al. Genome-wide screens identify Toxoplasma gondii determinants of parasite fitness in IFNγ-activated murine macrophages. Nat. Commun. 11, 5258 (2020).

48. Gold, D. A. et al. The Toxoplasma Dense Granule Proteins GRA17 and GRA23 Mediate the Movement of Small Molecules between the Host and the Parasitophorous Vacuole. Cell Host Microbe 17, 642–652 (2015).

49. Sangaré, L. O. et al. In Vivo CRISPR Screen Identifies TgWIP as a Toxoplasma Modulator of Dendritic Cell Migration. Cell Host Microbe 26, 478–492.e8 (2019).

50. Tachibana, Y., Hashizaki, E., Sasai, M. & Yamamoto, M. Host genetics highlights IFN-γ-dependent Toxoplasma genes encoding secreted and non-secreted virulence factors in in vivo CRISPR screens. Cell Rep. 42, 112592 (2023).

51. Giuliano, C. J. et al. CRISPR-based functional profiling of the Toxoplasma gondii genome during acute murine infection. Nat. Microbiol. 9, 2323–2343 (2024).

52. Krishnamurthy, S. et al. CRISPR Screens Identify Toxoplasma Genes That Determine Parasite Fitness in Interferon Gamma-Stimulated Human Cells. mBio 14, e00060–23 (2023).

53. Fox, B. A. et al. Toxoplasma gondii Parasitophorous Vacuole Membrane-Associated Dense Granule Proteins Orchestrate Chronic Infection and GRA12 Underpins Resistance to Host Gamma Interferon. mBio 10, e00589–19 (2019).

54. Ressurreição, M. et al. Use of a highly specific kinase inhibitor for rapid, simple and precise synchronization of Plasmodium falciparum and Plasmodium knowlesi asexual blood-stage parasites. PLoS ONE 15, e0235798 (2020).

55. Wang, J.-L. et al. Novel roles of dense granule protein 12 (GRA12) in Toxoplasma gondii infection. FASEB J. 34, 3165–3178 (2020).

56. Murillo-Léon, M., Bastidas-Quintero, A. M. & Steinfeldt, T. Decoding Toxoplasma gondii virulence: the mechanisms of IRG protein inactivation. Trends Parasitol. S1471–4922(24)00201–0 (2024) doi:10.1016/j.pt.2024.07.009.

57. Michelin, A. et al. GRA12, a Toxoplasma dense granule protein associated with the intravacuolar membranous nanotubular network. Int. J. Parasitol. 39, 299–306 (2009).

58. Etheridge, R. D. et al. The Toxoplasma pseudokinase ROP5 forms complexes with ROP18 and ROP17 kinases that synergize to control acute virulence in mice. Cell Host Microbe 15, 537–550 (2014).

59. Vella, S. A. et al. The role of potassium and host calcium signaling in Toxoplasma gondii egress. Cell Calcium 94, 102337 (2021).

60. Chen, Y., Li, X., Yang, M. & Liu, S.-B. Research progress on morphology and mechanism of programmed cell death. Cell Death Dis. 15, 1–13 (2024).

61. Coombs, R. S. et al. Immediate Interferon Gamma Induction Determines Murine Host Compatibility Differences between Toxoplasma gondii and Neospora caninum. Infect. Immun. 88, e00027–20 (2020).

62. Sokol, S. L. et al. Dissection of the in vitro developmental program of Hammondia hammondi reveals a link between stress sensitivity and life cycle flexibility in Toxoplasma gondii. eLife 7, e36491 (2018).

63. Donahoe, S. L., Lindsay, S. A., Krockenberger, M., Phalen, D. & Šlapeta, J. A review of neosporosis and pathologic findings of Neospora caninum infection in wildlife. Int. J. Parasitol. Parasites Wildl. 4, 216–238 (2015).

64. Cygan, A. M. et al. Proximity-Labeling Reveals Novel Host and Parasite Proteins at the Toxoplasma Parasitophorous Vacuole Membrane. mBio 12, e0026021 (2021).

65. Nyonda, M. A. et al. Toxoplasma gondii GRA60 is an effector protein that modulates host cell autonomous immunity and contributes to virulence. Cell. Microbiol. 23, e13278 (2021).

66. Wang, Y. et al. Three Toxoplasma gondii Dense Granule Proteins Are Required for Induction of Lewis Rat Macrophage Pyroptosis. mBio 10, e02388–18 (2019).

67. Donald, R. G., Carter, D., Ullman, B. & Roos, D. S. Insertional tagging, cloning, and expression of the Toxoplasma gondii hypoxanthine-xanthine-guanine phosphoribosyltransferase gene. Use as a selectable marker for stable transformation. J. Biol. Chem. 271, 14010–14019 (1996).

68. Huynh, M.-H. & Carruthers, V. B. Tagging of endogenous genes in a Toxoplasma gondii strain lacking Ku80. Eukaryot. Cell 8, 530–539 (2009).

69. Butterworth, S., et al. Mapping Host-Microbe Transcriptional Interactions by Dual Perturb-Seq. http://biorxiv.org/lookup/doi/10.1101/2023.04.21.537779 (2023) doi:10.1101/2023.04.21.537779.

70. Hunt, A. et al. Differential requirements for cyclase-associated protein (CAP) in actin-dependent processes of Toxoplasma gondii. eLife 8, e50598 (2019).

71. Robinson, J. T., Thorvaldsdóttir, H., Wenger, A. M., Zehir, A. & Mesirov, J. P. Variant Review with the Integrative Genomics Viewer. Cancer Res. 77, e31–e34 (2017).

72. Schindelin, J., et al. Fiji: an open-source platform for biological-image analysis. Nat. Methods 9, 676–682 (2012).

73. Alvarez-Jarreta, J. et al. VEuPathDB: the eukaryotic pathogen, vector and host bioinformatics resource center in 2023. Nucleic Acids Res. 52, D808–D816 (2024).

74. Jumper, J. et al. Highly accurate protein structure prediction with AlphaFold. Nature 596, 583–589 (2021).

