## Supplementary Figures and Legends for "*In vivo* CRISPR screens identify GRA12 as a transcendent secreted virulence factor across *Toxoplasma gondii* strains and mouse subspecies"

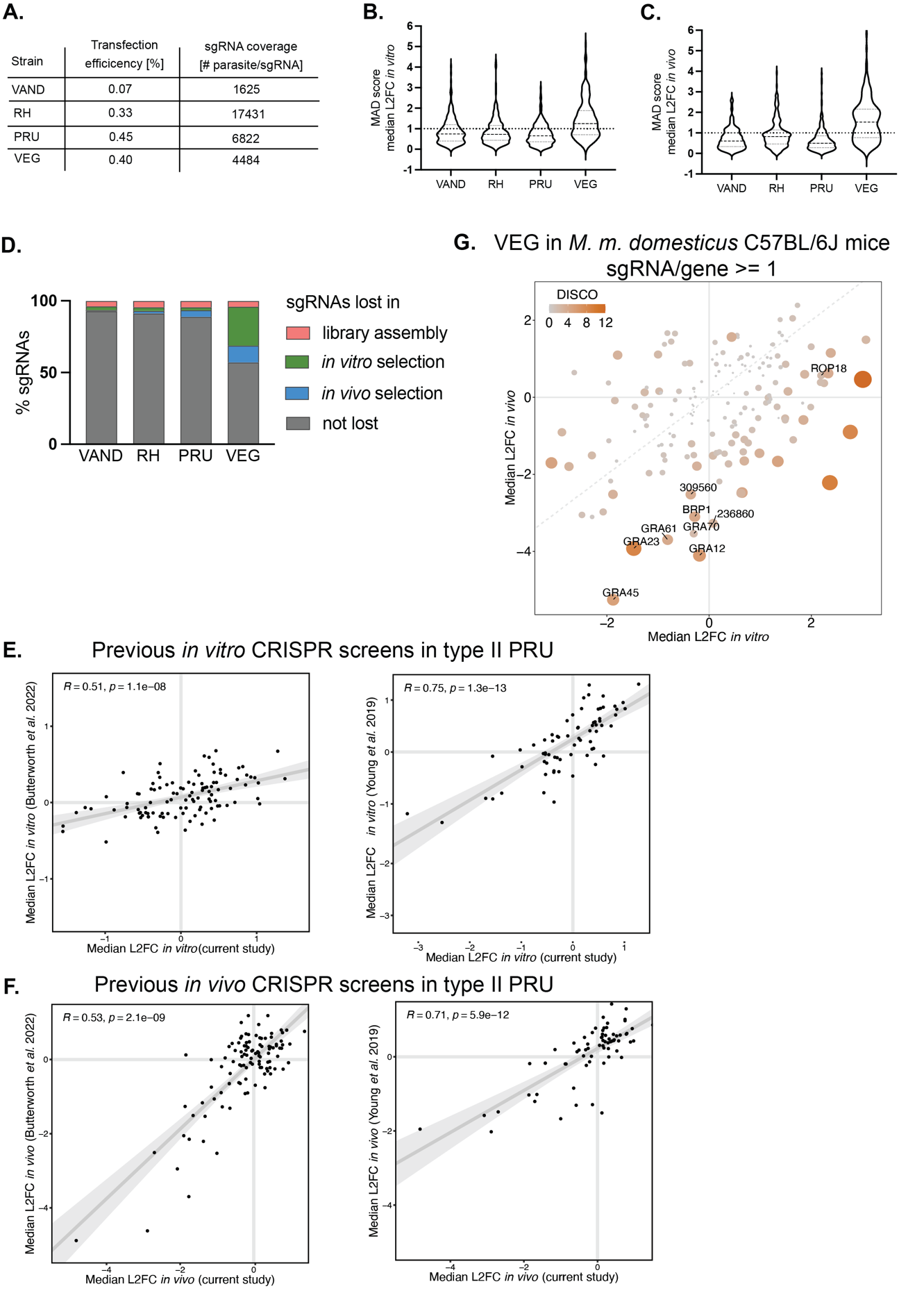

**Figure S1.** ***In vivo* CRISPR screens of the secretome of four *Toxoplasma* strains.** (A) Transfection efficiency and relative sgRNA coverage in each screen. (B) Median Absolute Deviation (MAD) of the log2 fold change (L2FC) of sgRNA read counts between the plasmid and the inoculum and (C) of the inoculum and the mouse peritoneum in each screen. (D) sgRNA loss at each screen step. (E) Correlation of the median L2FC *in vitro* and (F) *in vivo* of the current type II screen with previous screens performed in our research group ^42,43^. (G) Scatter plots of the median L2FC for each gene *in vitro* and *in vivo* of the CRISPR screen of VEG in C57BL/6J mice. The colour and size of each point reflects the Discordance/Concordance (DISCO) score, and the dashed grey line indicates equal L2FC. A threshold of 1 sgRNA per gene was applied compared to 3 sgRNAs per gene in Figure 1B.

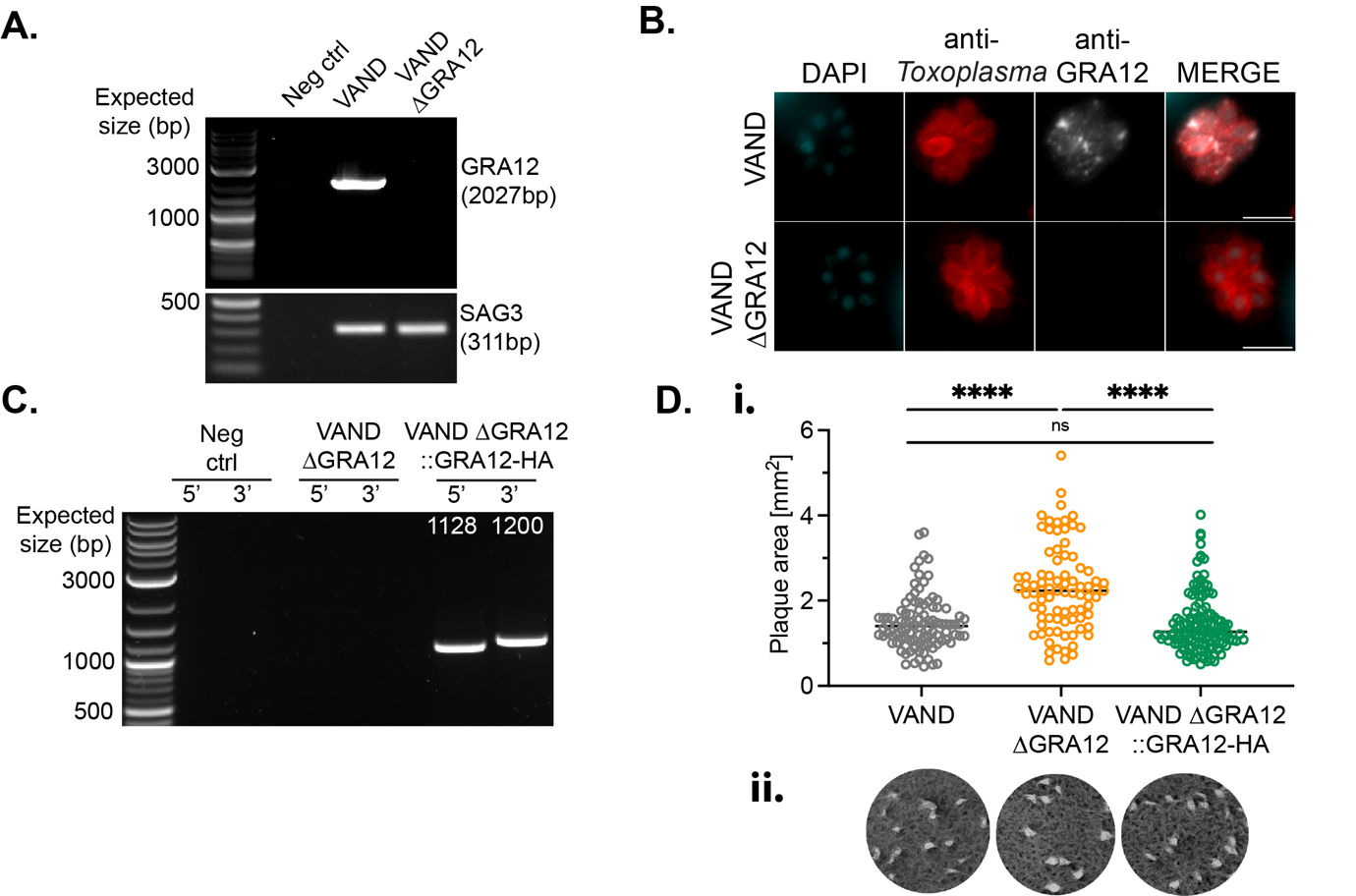

**Figure S2. Establishment of VAND ∆GRA12 and ∆GRA12::GRA12-HA strains.** (A) PCR amplification of the GRA12 locus in the parental VAND and ∆GRA12 strains. Amplification of SAG3 was performed as control. (B) Immunofluorescence verification of GRA12 in the VAND ∆GRA12 strain. Scale bar represents 10 µm. (C) PCR validation of the VAND ∆GRA12::GRA12-HA strain. (D) Scatter plot of the plaque size of VAND parental and derived clones, bar represents the median. Significance was tested using the one-way ANOVA test, N=2 (i) and respective images (ii).

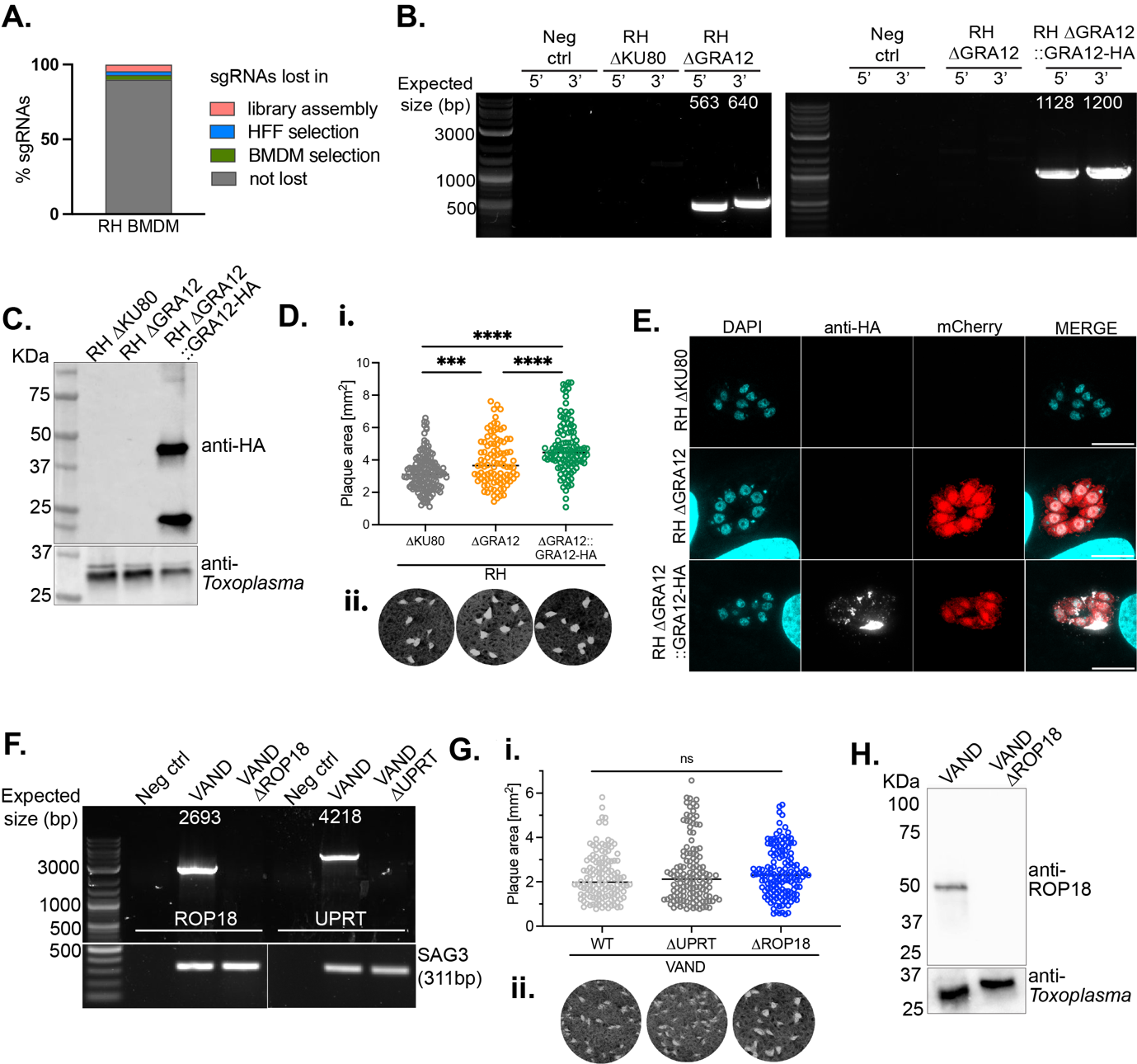

**Figure S3. Creation of RH and VAND KO mutants for the validation of GRA12 as transcendent virulence factor *in vitro*.** (A) sgRNA loss at each screen step of the type I RH CRISPR screen in BMDMs. (B) PCR validation of the RH ∆GRA12 and complemented strains. (C) Verification of the GRA12::HA expression via anti-HA detection in the RH ∆GRA12::GRA12-HA strain by western blot. (D) Plaque size of RH ∆KU80 parental and derived clones. Significance was tested using a one-way ANOVA, N=2, p *** <0.001, *** <0.0001 (i), and respective images (ii). (E) Immunofluorescence verification of the C-terminal HA-tagged GRA12 in the RH ∆GRA12::GRA12-HA strain. Scale bar is 10 µm. (F) PCR validation of the VAND ∆UPRT and ∆ROP18. Amplification of SAG3 was performed as control. (G) Plaque size of VAND parental and derived clones ∆UPRT and ∆ROP18. Significance was tested using an unpaired t test, N=1 (i), and respective images (ii). (H) Verification of the ROP18 KO via anti-ROP18 detection in the VAND ∆ROP18 strain by western blot.

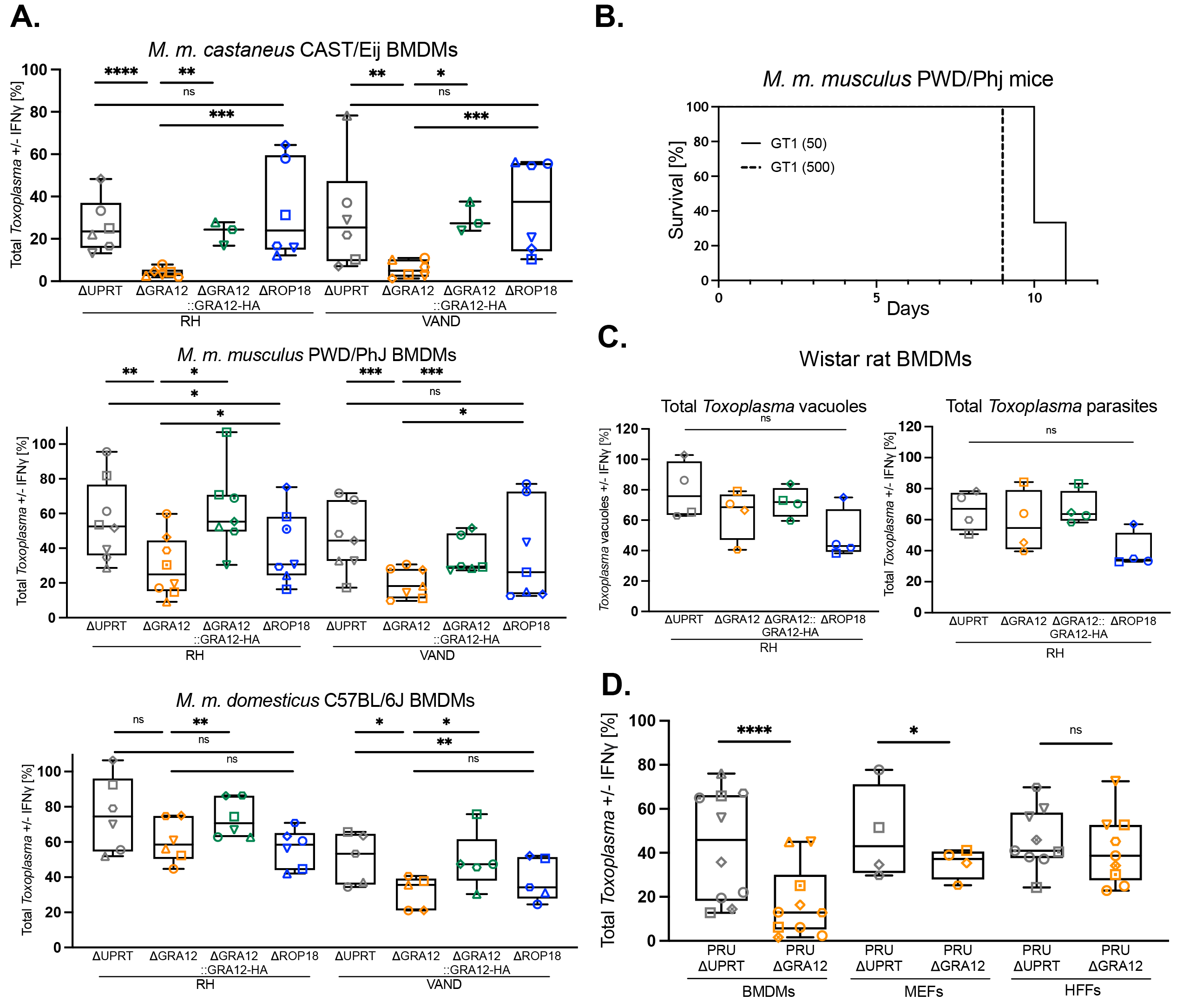

**Figure S4. Validation of GRA12 as *Toxoplasma* transcendent virulence factor in mouse subspecies.** (A) Quantification of high content-automated imaging of parasite numbers in IFNγ-treated BMDMs of different mouse subspecies relative to untreated controls. Cells were infected with RH ∆UPRT, ∆GRA12, ∆GRA12::GRA12-HA and ∆ROP18 and *Toxoplasma* parasites were quantified at 24 h after infection. (B) Survival curve of PWD/Phj mice infected with the type I GT1 strain, dose in parenthesis. N=3 mice per group. (C) Quantification of high-content automated imaging of *Toxoplasma* infection in IFNγ-treated Wistar rat BMDMs or (D) murine embryonic fibroblasts (MEFs), human foreskin fibroblasts (HFFs) and murine BMDMs, relative to untreated controls. Cells were infected with RH ∆UPRT, ∆GRA12, ∆GRA12::GRA12-HA and ∆ROP18, or PRU ∆UPRT and PRU ∆GRA12, and parasites were quantified at 24 h after infection. Symbol shapes indicate biological repeats. Significance was tested using the One-way Anova test with the Benjamini, Krieger and Yekutieli FDR correction. p * <0.05, ** <0.01, *** <0.001, **** <0.0001.

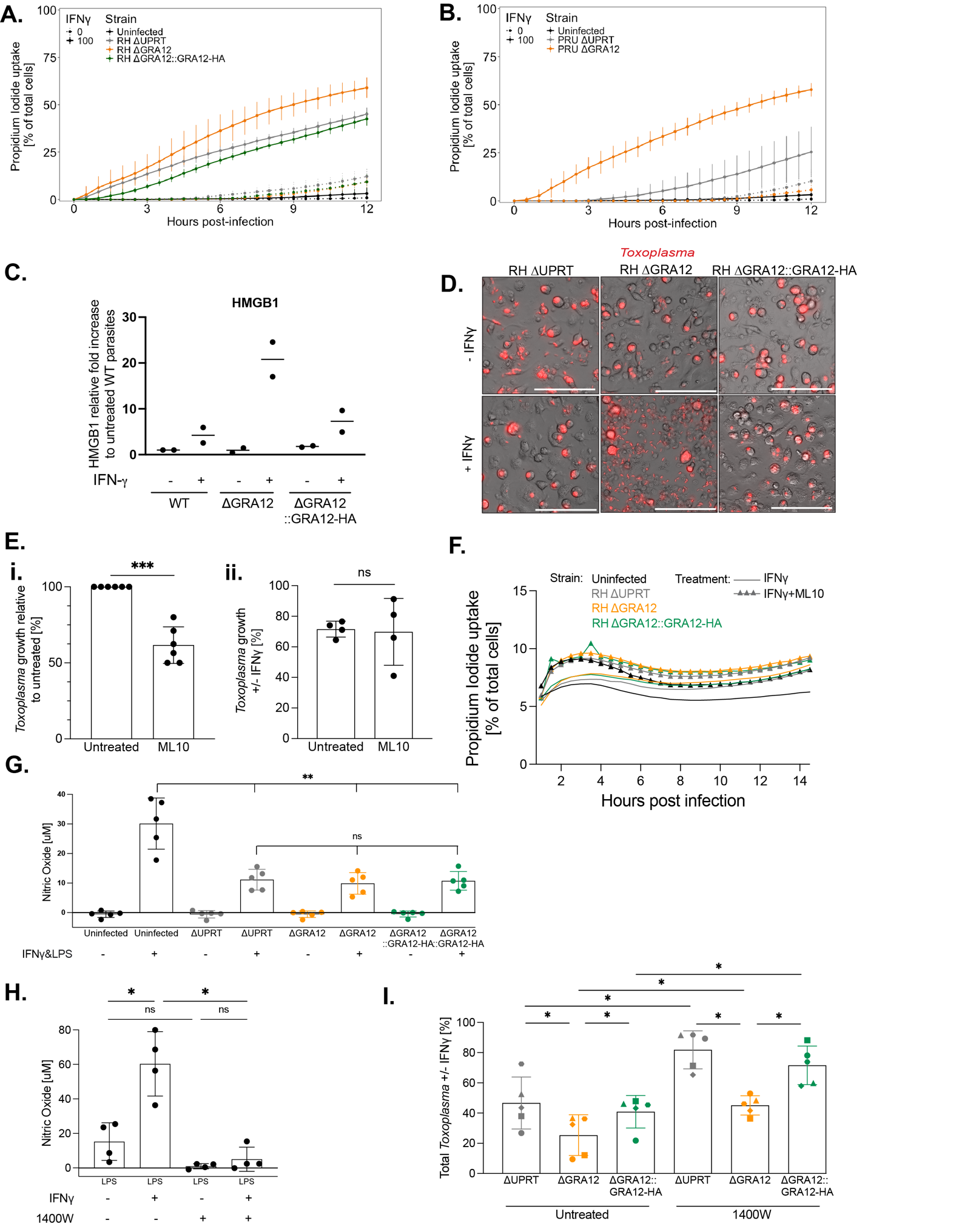

**Figure S5. Lack of GRA12 results in IFNγ-mediated host cell death by necrosis.** Time course of PWD/Phj BMDMs cell death quantified via Propidium Iodide uptake. Cells were pre-stimulated for 24 h with IFNγ (continue line) or left untreated (dashed line) and then infected with (A) RH ∆UPRT, ∆GRA12, ∆GRA12::GRA12-HA or (B) PRU ∆UPRT or ∆GRA12, or left uninfected as a control. The number of dead cells is expressed as a percentage of the total at each time point. (C) Quantification of the HMGB1 necrosis marker from the western blot analysis of the supernatant of infected BMDMs, relative to the untreated parental condition. (D) Microscopy images of IFNγ-treated or untreated C57BL/6J BMDMs, infected for 24 h with RH ∆UPRT, ∆GRA12 or ∆GRA12::GRA12-HA. Scale bar represents 100 µm. (E) Quantification by imaging of *Toxoplasma* growth in ML10-treated relative to untreated BMDMs, in naïve conditions (i) or in IFNγ-treated conditions relative to untreated controls (ii). (F) Time course of C57BL/6J BMDMs cell death quantified via Propidium Iodide uptake. Cells were infected with RH ∆UPRT, ∆GRA12, ∆GRA12::GRA12-HA or left uninfected, and 1 hpi treated with 1 µM ML10 (triangle symbol) or left untreated as a control (continue line). The number of dead cells is expressed as a percentage of the total at each time point. (G) Quantification of Nitric Oxide from C57BL/6J BMDMs stimulated for 24 h with 100 U/ml IFNγ and 0.2 µg/ml LPS or left untreated as control, and infected for 24 h with RH ∆UPRT, ∆GRA12, ∆GRA12::GRA12-HA or left uninfected. Significance was tested using the One-way Anova test with the Benjamini, Krieger and Yekutieli FDR correction, (H) Quantification of Nitric Oxide from a culture of uninfected C57BL/6J BMDMs pretreated for 24 h with 100 U/ml IFNγ and with the iNOS inhibitor 1400W, or left untreated as control, and stimulated for 24 h with 0.2 µg/ml LPS. Significance was tested using the One-way Anova test with the Benjamini, Krieger and Yekutieli FDR correction, (I) Quantification by imaging of *Toxoplasma* restriction in IFNγ-treated or untreated C57BL/6 BMDMs, pretreated of not with 1400W, infected with RH ∆UPRT, ∆GRA12, ∆GRA12::GRA12-HA and parasites were quantified at 24 h after infection. Symbol shapes indicate biological repeats. P *<0.05, ** < 0.01.

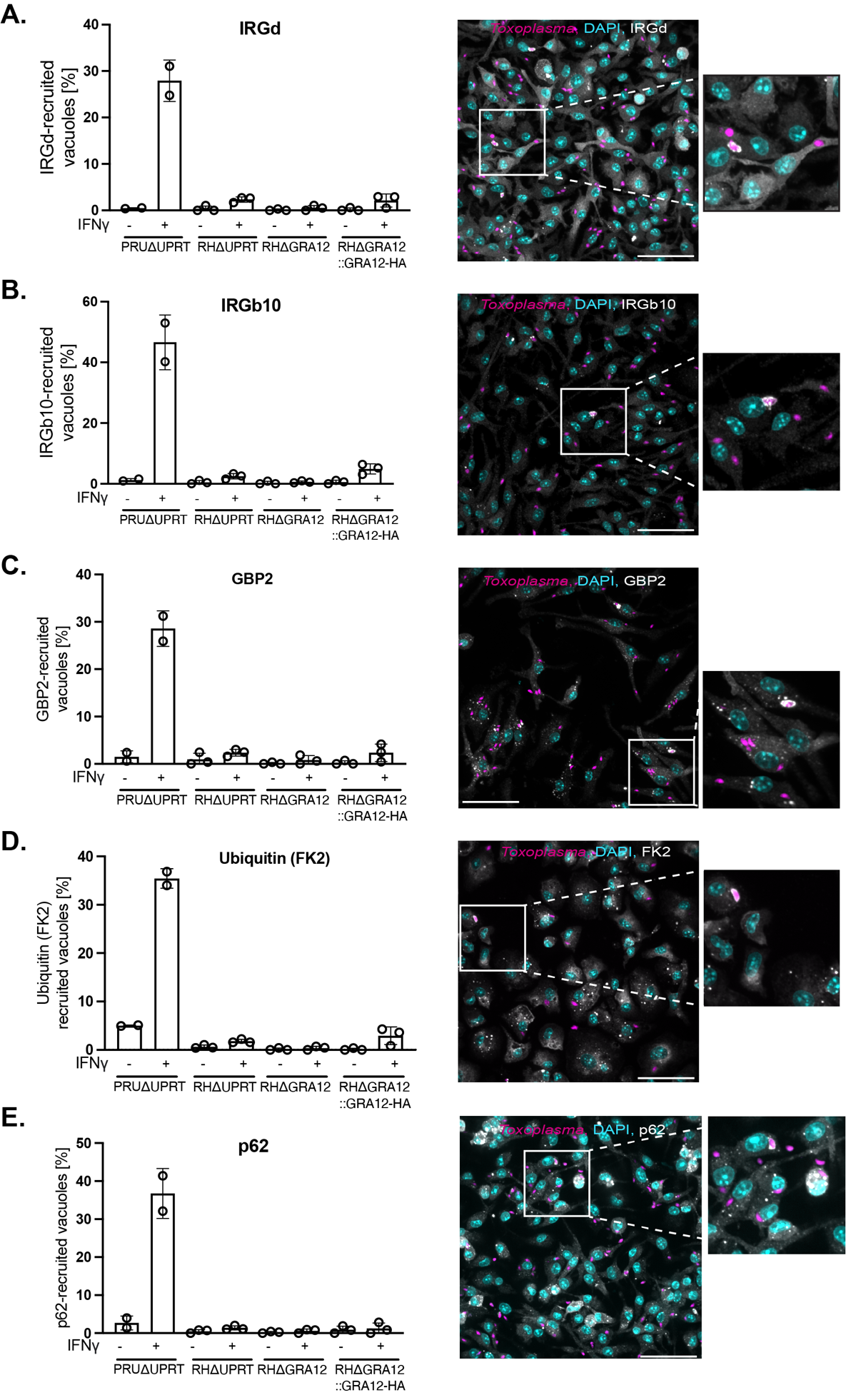

**Figure S6. Recruitment of host factors to the PVM.** Quantification of the recruitment to the PVM of host proteins: IRGd (A), IRGb10 (B), GBP2 (C), Ubiquitin (FK2, D) and p62 (E) with representative images on the right panels. C57BL/6J BMDMs were pretreated for 24h with IFNγ or left untreated, and infected for 90 minutes with RH ∆UPRT, ∆GRA12 or ∆GRA12::GRA12-HA, before being fixed and stained for immunofluorescence. N=3 and N=2 for the PRU ΔUPRT line used as positive control. Scale bar represents 50 µm.

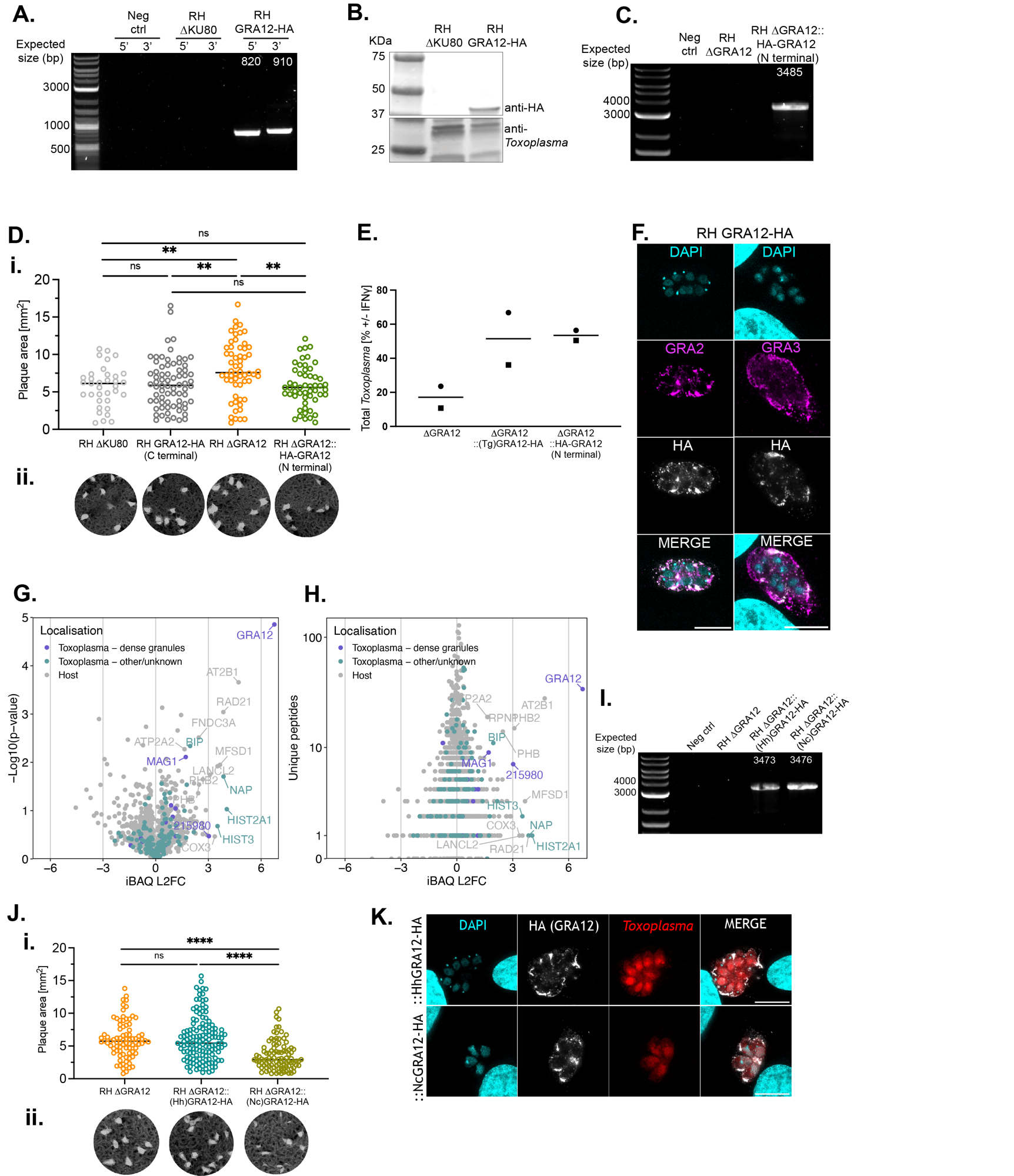

**Figure S7. Creation of the N and C terminally HA-tagged GRA12 strains and validation of GRA12 paralogs activity.** (A) PCR validation of the RH GRA12 C-terminally HA-tagged line in the endogenous locus (RH GRA12-HA) by PCR and (B) by western blot. (C) PCR validation of the RH GRA12 N-terminally HA-tagged line (RH ∆GRA12::HA-GRA12) created by complementing GRA12 in the *Uprt* locus. (D) Plaque size of RH GRA12-HA and RH ∆GRA12::HA-GRA12 lines compared to their respective parental RH ∆KU80 and RH ∆GRA12 strains. Significance was tested using an unpaired t test, N=1 (i) and respective images (ii). (E) Relative *Toxoplasma* growth in IFNγ-treated versus untreated BMDMs. BMDMs were infected with the RH ∆GRA12 strain, or the RH∆GRA12 strain complemented with *Toxoplasma* (Tg) GRA12 or the N-terminally HA-tagged line RH ∆GRA12::HA-GRA12, or the parental RH ∆GRA12 strain as control for 24h before a plate reader quantification of the mCherry signal as proxy for parasite growth. N=2. (F) Immunofluorescence localisation of GRA2 (left panel) and GRA3 (right panel) in relation to GRA12 in the RH GRA12-HA line infecting HFFs. Scale bar represents 10 µm. (G) Volcano plot of proteins identified by mass spectrometry in the anti-HA pull down in lysates from IFNγ-treated BMDM infected with either RH GRA12-HA or the parental RH ∆KU80 strains. The relative abundance of the proteins (iBAQ L2FC) is plotted against the p-value (-Log10(p-value), (G)) and against the number of peptides detected ((Unique peptides), (H)). (I) PCR validation of the RH ∆GRA12 strain complemented with the *H. hammondi* (::(Hh)GRA12-HA) or the *N. caninum* (::(Nc)GRA12-HA) GRA12 homologues. (J) Plaque size of the complemented ::(Hh)GRA12-HA and ::(Nc)GRA12-HA strains, compared to the parental RH ∆GRA12 strain. Significance was tested using an unpaired t test, N=1 (i) and respective images (ii). (K) Immunofluorescence localisation of GRA12 via anti-HA in the RH ∆GRA12::(Hh)GRA12-HA strain (upper panel) and RH ∆GRA12::(Nc)GRA12-HA strain (lower panel). Scale bar represents 10 µm. p ** <0.01, **** <0.0001.

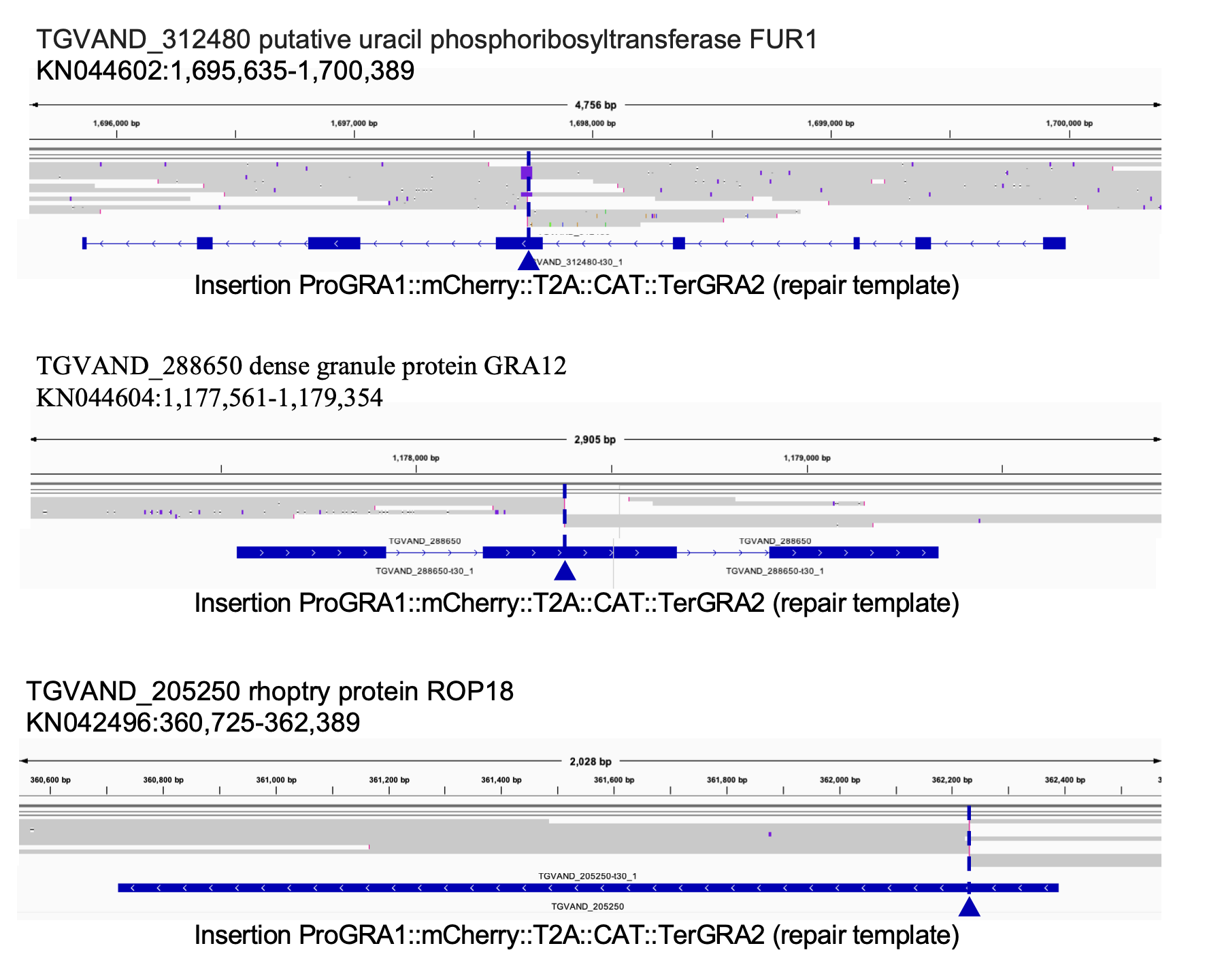

**Figure S8. Validation by Nanopore sequencing of the VAND mutant strains.** Alignment of reads from Nanopore sequencing to the reference genome assembly ToxoDB-65_TgondiVAND. The alignment shows disruption of the endogenous UPRT (TGVAND_312480), GRA12 (TGVAND_205250) and ROP18 (TGVAND_205250) loci, for the creation of the VAND ΔUPRT, ΔGRA12 and ΔROP18 strains respectively. The arrowhead indicates the integration of the repair template ProGRA1::mCherry::T2A::CAT::TerGRA2 in the coding sequence.
